# Abundant Recurrent Mitochondrial Mutations and Widespread Mitonuclear Epistasis in *Caenorhabditis elegans*

**DOI:** 10.1101/2025.04.22.650036

**Authors:** Tuc H.M. Nguyen, Daniel A. Klein, Olivia S. Weklar, Erin R. Wengrow, Matthew V. Rockman

## Abstract

Coordinated genetic and physical interactions between mitochondrial and nuclear gene products regulate ATP production in the mitochondria. Linking mitochondrial genotypes and mitonuclear genetic interactions to phenotypes remains a complex challenge. Here, we have developed *Caenorhabditis elegans* as a model for mitonuclear epistasis studies. In a sample of 540 genetically distinct wild isolates, 10% of sites in the mitochondrial genome vary, with hundreds of missense mutations segregating in the species. Recurrent mutations and triallelic sites are common. Phylogenetic analyses of mitogenome sequences identified eight distinct lineages, each with diagnostic variants. Principal component analysis of the nuclear genomes showed considerable concordance between mitochondrial and nuclear genomes in *C. elegans* populations, suggesting that disrupting coevolved mitonuclear genetic combinations could reveal substantial epistasis. We used GPR-1 overexpression, which disrupts the first mitotic division, to efficiently exchange nuclear and mitochondrial genomes between all pairs of 18 naturally isolated *C. elegans* strains, generating the largest-to-date animal mitonuclear exchange panel, with 323 unique viable mitonuclear genotypes. We phenotyped development of a subset of strains, with 30 unique genotypes, under six different environmental conditions, including high temperature and exposure to heavy metals. Mitonuclear epistasis contributed significantly to phenotypic variance across all tested conditions. We also tested for mitonuclear coadaptation by comparing the stress resistance of matched and mismatched cybrids. Interestingly, some mismatched strains exhibited greater resistance, highlighting the complexity and context dependence of mitonuclear interactions.

## Introduction

Metazoan life hinges on a continuous supply of energy from the mitochondria, which governs virtually all cellular functions (Ponka 1999; Susin, et al. 1999; Duchen 2000; Mandal, et al. 2011). Both mitochondrial and nuclear genomes contribute to mitochondrial respiration, as oxidative phosphorylation enzyme complexes are composed of subunits from both genomes (Huynen, et al. 2013; van der Sluis, et al. 2015; Elurbe and Huynen 2016; Sloan, et al. 2018). Changes in either genome could disrupt this balance, impacting not only direct protein-protein interactions (Ellison and Burton 2006; Osada and Akashi 2012; Havird, Whitehill, et al. 2015), but also the broader mitochondrial processes (Chou, et al. 2010; Ellison and Burton 2010; Meiklejohn, et al. 2013). The retrograde-anterograde signaling pathways between the nucleus and the mitochondria represent another arena for complex mitonuclear interactions that are extremely sensitive to environmental changes (Wallace 2012; Baris, et al. 2017; Weinhouse 2017).

To maintain functional mitochondria and organismal fitness, natural selection should favor coadapted combinations of mitochondrial and nuclear alleles. Hybridization can disrupt coevolved and coadapted combinations of mitochondrial and nuclear genomes, leading to hybrid breakdown due to Dobzhansky-Muller incompatibilities (DMI) (Rand, et al. 2004; Sloan, et al. 2017). Interspecific studies, such as those in *Drosophila* (Montooth, et al. 2010; Hoekstra, et al. 2013; Meiklejohn, et al. 2013), *Saccharomyces* (Zeyl, et al. 2005; Lee, et al. 2008; Chou, et al. 2010; Spirek, et al. 2014; Jhuang, et al. 2017), and *Xiphophorus* fishes (Moran, et al. 2024), have been key to understanding these incompatibilities across millions of years of divergence. In contrast, studying intraspecific mitonuclear interactions in diverging populations offers the opportunity to capture the gradual emergence of mitonuclear DMIs, where subtle mutations and compensatory changes evolve to maintain mitochondrial functions, long before complete reproductive isolation occurs. This approach offers an effective model for understanding the early stages of mitonuclear DMIs, and how these interactions contribute to population structure and speciation over shorter evolutionary timescales.

Introducing distinct mtDNAs into multiple different nuclear backgrounds is a powerful strategy for detecting mitonuclear epistasis within a species, including budding yeast (Paliwal, et al. 2014; Wolters, et al. 2018; Nguyen, et al. 2020; Biot-Pelletier, et al. 2023; Nguyen, et al. 2023), fruit flies (Rand, et al. 2018; Mossman, Ge, et al. 2019; Vaught, et al. 2020), and copepods (Ellison and Burton 2008; Burton and Barreto 2012; Burton, et al. 2013). In animals, the standard experimental design to isolate the effects of mitonuclear interactions has been backcrossing for multiple generations (Rand, et al. 2018; Mossman, Biancani, et al. 2019; Mossman, Ge, et al. 2019; Camus, et al. 2020). However, during the breeding process, selection can alter allele frequencies in unpredictable ways, which can confound interpretation of experimental results, and allow new mutations to accumulate in the final strains. Budding yeast offers the ability to exchange mitochondrial DNA in a single genetic step, and access to abundant genomic resources resulting from global sequencing efforts (Peter, et al. 2018; De Chiara, et al. 2020). Studies of yeast, however, are limited to questions related to cellular and molecular phenotypes, but not the life-history traits unique to metazoans, such as fecundity, developmental rates, and behavioral phenotypes.

The genetically tractable and easily manipulated nematode *Caenorhabditis elegans* emerges as a highly promising model for the investigation of mitonuclear coadaptation. The recent development of population-genomic resources for *C*. *elegans* wild isolates (Cook, et al. 2017; Crombie, et al. 2019; Lee, et al. 2021) opens extensive opportunities to explore previously untapped genetic variation in the mitochondrial genomes. Given the presence of population structure within *C. elegans* (Lee, et al. 2021), and the uniparental inheritance and high mutation rate of mtDNA (Konrad, et al. 2017), it is reasonable to anticipate considerable divergence in mitochondrial genomes among populations. Although selfing has led to an overall reduction in genetic diversity relative to outcrossing species (Graustein, et al. 2002; Cutter and Payseur 2003), *C. elegans* has paradoxically maintained hyperdivergent genomic regions harboring tremendous diversity, comparable to that found between sister species (Lee, et al. 2021). This suggests that exchanging mtDNAs between individuals from distinct populations should yield novel genetic combinations and, consequently, novel phenotypic outcomes.

It is now possible to precisely exchange nuclear and mitochondrial genomes between strains in *C. elegans* in a two-step experimental process taking advantage of the GPR-1 protein, which regulates chromosomal segregation in stages leading to the first cleavage in the *C. elegans* embryos (Artiles, et al. 2019). During the first mitotic prophase, overexpression of the GPR-1 protein leads to premature separation of the maternal and paternal pronuclei, preventing proper mixing and segregation of parental chromatids into the first two blastomeres (AB and P1) (Besseling and Bringmann 2016). This event produces viable chimeric worms, with cells derived from AB lineage homozygous for maternal chromosomes and cells derived from the P1 lineage — including the germline — homozygous for paternal chromosomes. Such chimeric worms upon self-fertilization create offspring that contain a nuclear genome entirely derived from the paternal parent, and a maternally inherited mitochondrial genome (Zhou, et al. 2011).

Here, we have established *Caenorhabditis elegans* as a model organism for the investigation of mitonuclear interactions. We analyzed the population structure of mitochondrial genomes in *C. elegans* natural populations and revealed a wealth of polymorphisms spanning the entire genome, including its protein coding regions. Furthermore, we detected considerable genomic concordance between the nuclear and mitochondrial genome. Leveraging the GPR-1 overexpression construct, we engineered a comprehensive collection of mitonuclear cytoplasmic hybrid (cybrid) strains, amounting to an 18×18 grid consisting of 323 distinct viable genotypes, each a unique mitonuclear genetic combination. Subsequently, we systematically assessed phenotypes of a subset of these cybrids under varying mitochondrially taxing conditions, including exposure to paraquat, heavy metals, and high temperature. Our findings underscore the prevalence of mitonuclear epistasis across the examined conditions, dwarfing the main effects of mtDNA. Our results support the idea that the mitochondrial genome affects phenotypic variation at a far greater level than expected given its small size, while the pattern of effects provides surprisingly limited support for models of mitonuclear coadaptation. This work will help us disentangle genotype-phenotype links and provide a framework for better understanding the evolutionary processes that shape population structures and local adaptation.

## Results

### Mitochondrial phylogeny

We examined mitochondrial variants in the genomes of the 1385 wild isolates in the *Caenorhabditis* Nucleotide Diversity Resource, CaeNDR (Cook, et al. 2017; Crombie, et al. 2019; Lee, et al. 2021). This collection of strains includes many sets of extremely similar strains, often collected from single patches of substrate and likely to represent very close relatives descended by selfing from a single shared ancestor within a few generations. To reduce the influence of these related strains on population-genetic analysis, we focused on a reduced set of 550 isolates, defined in CaeNDR as isotype reference strains (Crombie, et al. 2019), each different in the nuclear genome from each other by at least 0.03% (we revisit the remaining strains for functional annotations below). Ten isotype reference strains with heterozygosity level greater than 10% for were removed from further analyses. The remaining 540 isotype reference strains included 367 unique mitochondrial haplotypes, with 1,458 segregating sites, of which 605 are singletons. While most mitochondrial haplotypes (324/367) are found only in a single isotype, the MY1 haplotype (found in 61 isotypes) and N2 haplotype (16 isotypes) were common, with MY1 hailing from every continent where *C. elegans* has been found — all but Antarctica (**Table S1**). Nuclear nucleotide diversity within the most common mitochondrial haplotypes, MY1 (π = 0.0018) and N2 (π = 0.0015), is lower than the population-wide diversity (π = 0.0033, N = 540), suggesting that these isotypes are relatively homogeneous.

To understand the extent to which recombination contributed to the genetic diversity of the mitochondrial genomes, we first performed recombination analysis on the whole genome alignment using RDP5 programs (Martin, et al. 2021). Consistent with the lack of recombination in the evolutionary history of metazoan mtDNA (Rokas, et al. 2003), we found no evidence for recombination across 6 different methods (**Methods**). Without recombination, the mitochondrial genome represents a single historical genealogy that we can reconstruct using phylogenetic inference. The maximum-likelihood tree for the complete mitogenome includes long internal branches, allowing for classification of the mitochondrial genomes into 8 distinct phylogenetic groups (**Figure 1A**). Hawaiian isolates were dispersed across the tree, consistent with the findings that Hawaiian strains harbor the most genetic diversity in the *C. elegans* populations (Crombie, et al. 2019; Lee, et al. 2021). Nevertheless, haplogroups showed signatures of biogeographic history (**Table S1**). Haplogroups E, F, G, and H were predominantly found in Hawaiian strains. Haplogroup C consisted mainly of strains from the Atlantic coast of Europe and Atlantic islands (Madeira, Sao Tome, and the Azores). Most of the 540 isotype reference strains carry mitochondrial haplotypes from Haplogroups A and B, which harbored strains predominantly from Europe, Australia, and the Americas. Notably, the genome reference strains N2 and CB4856 both carried haplotypes from Haplogroup B. Most interestingly, the mitochondrial genome of strain ECA1298 diverged significantly from all other mitochondrial genomes, forming its own distinct haplogroup. It differed by 108 mutations from its closest mitochondrial relative. The ECA1298 isotype is known only from Kalopa State Recreation Area (Big Island, Hawaii), where it has been collected several times.

**Figure 1.**
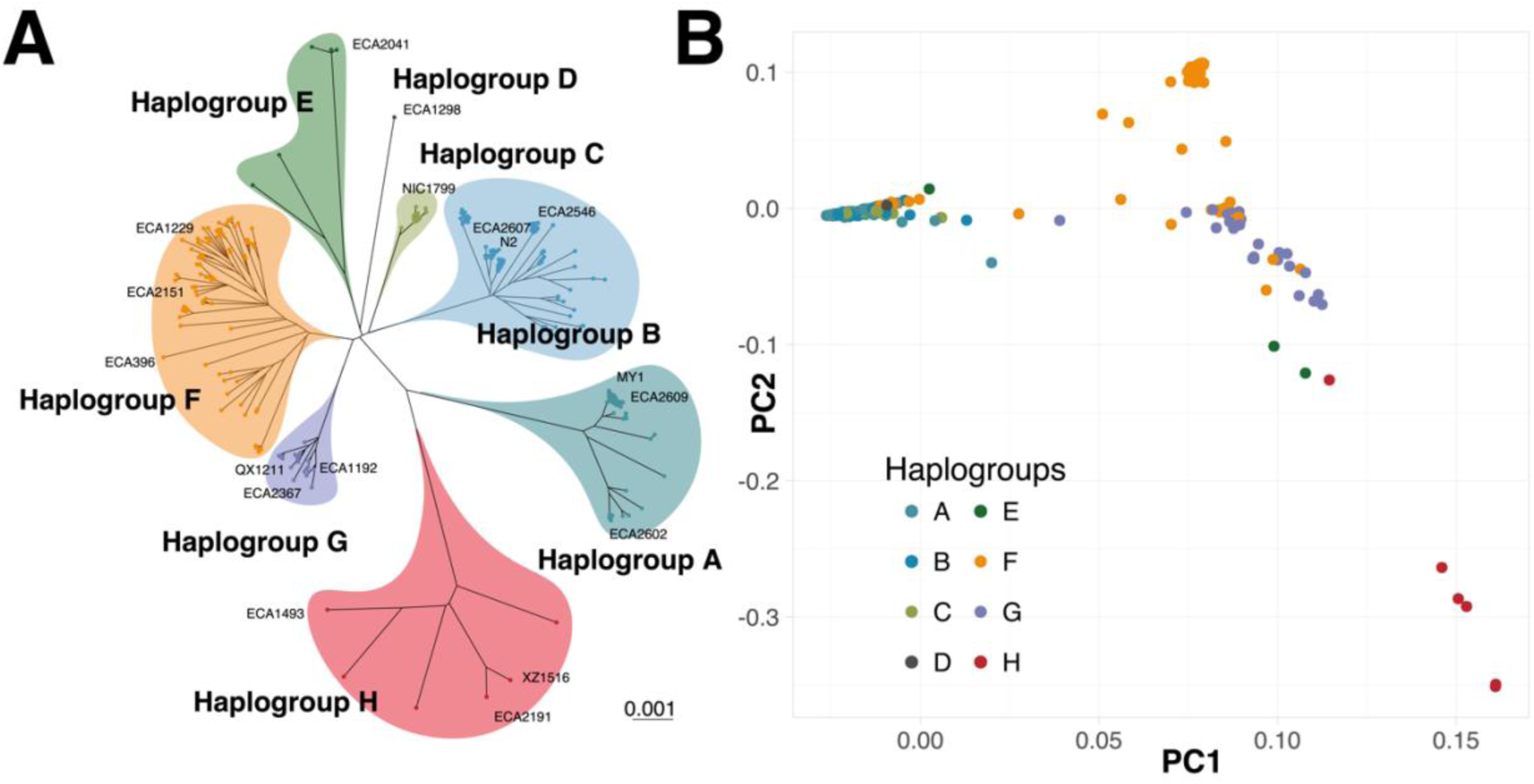
Mitochondrial population structure of *C. elegans*. **A. Maximum Likelihood Phylogenetic tree** of 367 unique mtDNA sequences of wild isolates, displaying 8 distinct mitochondrial lineages. Strains used in the subsequent reciprocal mitochondrial swap experiment are labeled. **B. Principal Component Analysis of Nuclear SNP genotypes, with each strain colored by mitochondrial haplogroup,** shows mitonuclear concordance in *C. elegans* natural populations.

To better understand the evolutionary history of *C. elegans*, we assembled the mitochondrial genome of its sister species, *C. inopinata* (Kanzaki, et al. 2018), and used it as an outgroup. We estimated the synonymous divergence (dS = 6.72) and the non-synonymous divergence (dN = 0.34) between the species’ mitochondrial genomes. When *C. inopinata* was included in the phylogenetic analysis with the *C. elegans* mitochondrial strains, it suggested that the root of the *C. elegans* tree is nested within Haplogroup E (**Figure S1**). However, this rooting implies that some isolates from Haplogroup E have remained nearly unchanged since the origin of the species, while others have undergone substantial evolutionary changes. We found it more plausible that the branch connecting *C. inopinata* to the *C. elegans* tree is saturated with substitutions at evolvable sites, obscuring evolutionary signals and making it challenging to accurately polarize mutations within *C. elegans*. Therefore, *C. inopinata* may not be suitable for accurately determining the location of the *C. elegans* mitochondrial root.

### Functional genetic variation in *C. elegans* mitochondrial genomes

*C. elegans* mitochondrial DNA content and organization consists of 12 protein-coding genes, two ribosomal RNAs, 22 tRNAs, and an AT-rich non-coding region (D-loop) (Okimoto, et al. 1992; Lemire 2005). Analysis of these genes revealed multiple segregating variants affecting protein sequences, tRNAs, and rRNAs (**Table S2**). We found missense variants in every protein-coding gene, with 8.4% of all codons (289/3433) showing missense variants within our sample. Absent tools for precise genetic engineering or recombination of mitochondrial genomes, options to experimentally assess the effects of single mitochondrial mutations are limited. We examined the 367 unique mitochondrial haplotypes among the 540 isotype strains to identify haplotype pairs that differ by a single mutation (**Table S3)**; contrasts between strains that differ only at these mitochondrial loci then reveal the effects of single mitochondrial mutations. We found 144 mutations, including 63 missense and 26 RNA mutations, that can be tested individually. Restricting the analysis to single missense differences with the N2 reference genome, there are 6 individually accessible mutations in 6 different ORFs.

Additionally, we identified 28 mitochondrial variants and one in-frame indel that segregate among 832 co-isotypic strains (**Table S4**). As a result, these variants were not included in the 540-strain analyses. These intra-isotype variants include 19 missense variants, each of which can be studied in isolation by comparing strains that differ only at one such site.

We next tested whether the mitochondrial haplogroups are likely to represent functionally distinct genomes. Holding the mitochondrial tree constant, we inferred the minimum number of mutational changes per branch for each of the 311 missense and 161 rRNA and tRNA variants (i.e., ignoring synonymous and intergenic variants). For these potentially functional variants, our branch specific analyses revealed mutation counts ranging from 0 to 10 across all branches, with internal branches that connect all the mitochondrial haplogroups accumulating 35 mutations (**Figure S2A**). The long branch leading to ECA1298 mitochondrial haplotype carries 5 distinct mutations (**Figure S2A**, **Table S6**). In contrast, most of the mutations (563) were located at the tips of the branches, consistent with their having arisen recently and being subject to purifying selection (**Figure S2B-O**). Overall, 367 out of 472 potentially functional variants could be explained by a single mutational event on the tree, while 86 sites required recurrent mutations. The remaining 19 sites containing missing genotypes were not included in this analysis. Recurrent mutations were found in all protein coding genes except *ctc-3*, as well as in RNA-coding sequences (**Table S5).** Notably, the *nduo-1* gene had the highest recurrence, with 10 unique sites experiencing recurrent mutations, including MtDNA:2042, which underwent at least 14 independent changes (**Figure S3A**). Regardless, each of the haplogroups carries diagnostic functional mutations (**Figure S2A**, **Table S6**), including three recurrent mutations shared across multiple haplotypes: an 18s rRNA variant (MtDNA:911) in haplogroups A, E, and F; a missense mutation in *ctc-1* (MtDNA:9242) in haplogroups A and G; and a variant in tRNA-Leu (MtDNA:5668) in haplogroups H and F. Excluding these, there are 26 unique functional mutations distinguishing the haplogroups (**Table S6**).

Recurrent mutations in the mitochondrial genome violate the infinite sites assumption, limiting the applicability of many standard population genetic statistics in assessing evolutionary forces shaping mitochondrial genome evolution. To navigate this complexity, we adopted a phylogenetic approach using a suite of tools available within HyPhy (Kosakovsky Pond, et al. 2020). To isolate the influence of segregating deleterious mutations affecting dN/dS estimates, we separately considered internal and tip branches of the tree (Hahn, et al. 2002; Lorenzo-Redondo, et al. 2016). Given the presence of recurrent mutations and strong evidence for rate heterogeneity, we anticipated variable selection pressure at different sites across different branches of the phylogeny. Thus, we applied the Mixed Effects Model of Evolution (MEME) method to detect selection affecting only a subset of the phylogeny while maintaining a constant mitochondrial phylogeny (Murrell, et al. 2012). Under a full codon model, global dN/dS ratios for each gene ranged from 0.013 to 0.142 for the internal branches, and 0.042 to 0.383 for the tips of the phylogeny (**Table S7**). For all genes, internal branches have substantially lower dN/dS ratio than tip branches (interior/tip ratios range from 11% to 46% across genes, median 22%), suggesting that low-frequency nonsynonymous mutations on the tip branches may represent recent deleterious mutations, and that similar mutations have been purged over the longer time periods represented by internal branches (**Figure S2B-O)**. Interestingly, among all 12 protein-coding genes analyzed, MEME detected significant episodic diversifying selection in only one site at the FDR of 5%: the 94^th^ codon in *nduo-1* (*p* = 0.018), which corresponds to the most frequent recurrent variant MtDNA:2042, suggesting potential adaptive evolution at this site. Our results highlight pervasive purifying selection within the mitochondrial genome and suggest the potential for mitonuclear co-evolutionary dynamics, driven by adaptive changes at specific sites.

### Mitonuclear population structure

We next investigated the concordance of mitochondrial and nuclear genomes in *C. elegans* populations using principal component analysis of the nuclear genomes (**Figure 1B**). Nuclear PC1 largely separates the cosmopolitan, Atlantic, and Hawaiian strains carrying haplogroups A, B, D, and E from the primarily Hawaiian strains carrying haplogroups F, G, and H. PC2 in turn separates Hawaiian strains in a pattern largely concordant with mitochondrial haplogroup membership; for example, strains from mitochondrial haplogroup F largely occupy a separate region of nuclear PC2 space from strains with mitochondrial haplogroup H. Notwithstanding this general pattern, haplogroup G strains are distributed across much of the space of these first two nuclear PCs. Of the 216 isotypes that share identical mtDNA, 177 belong to haplogroups A and B (**Table S1**), whose nuclear genomes are tightly clustered in the PCA (**Figure 1B**). Overall, we see considerable concordance between mitochondrial and nuclear genomes from the PCA, suggesting that exchanging mtDNA between strains of different mitochondrial lineages could potentially disrupt mitonuclear genetic coadaptation and reveal substantial epistatic interactions.

### Construction of a mitonuclear panel

To generate strains with unique mitonuclear genotypes, we transferred mtDNAs among a subset of divergent isolates from the 540 wild worm strains. We employed GPR-1 overexpression to construct strains with unique mitonuclear combinations (18 nuclear x 18 mitochondrial). The creation of mitochondrial cybrid strains followed a two-step process (**Figure 2A**). Step 1 involved the transfer the mtDNAs of interest from original strain into a “*vehicle*” strain that carries a GPR-1 overexpression cassette and fluorescent markers allowing for quick identification of chimeric worms in cross progeny. Subsequently, the mtDNA from the *vehicle* strain was introduced into all 18 nuclear backgrounds (**Methods**).

**Figure 2.**
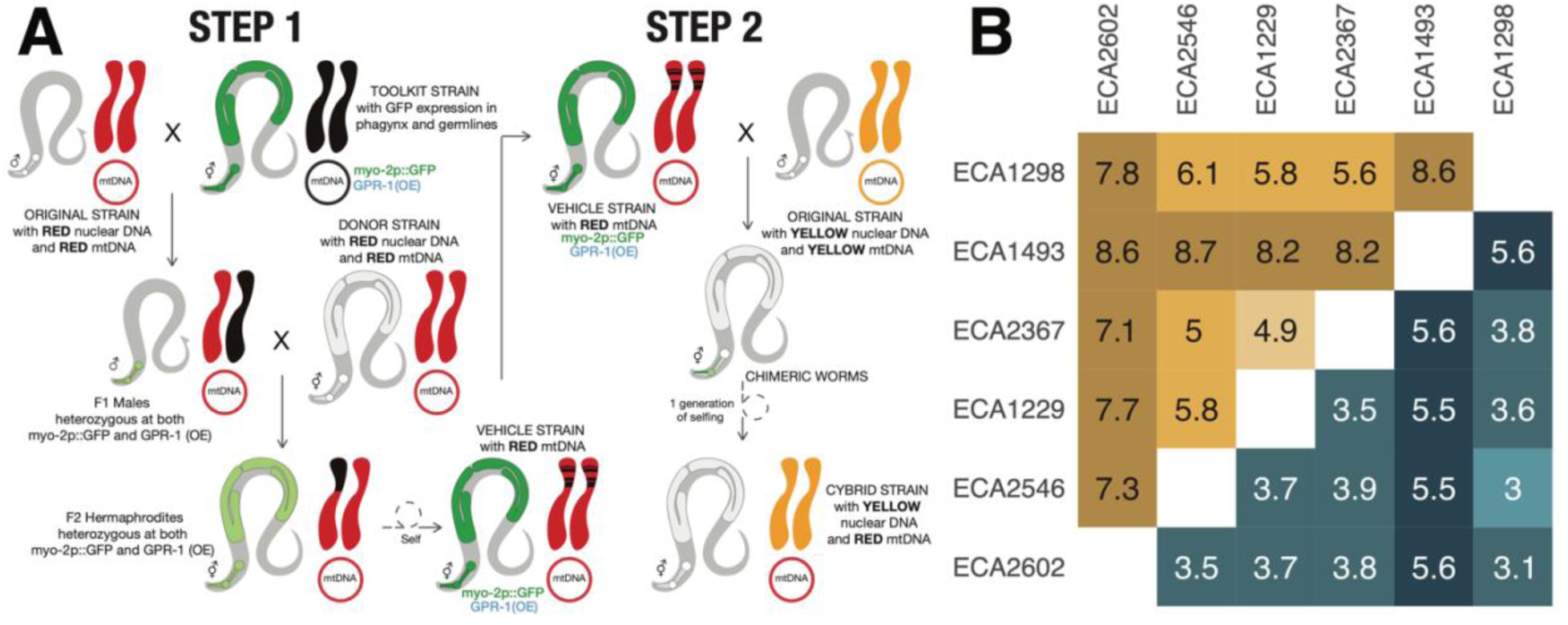
A. **Two-step scheme to exchange mitochondrial genome between *C. elegans* wild isolates.** Step 1 - constructing vehicle strains with different mtDNAs. Each of the parental males is mated to the GPR-1(OE) toolkit strain, which expresses GFP in pharyngeal tissues and germline. F_1_ heterozygous males are crossed to the wild strain hermaphrodite harboring the mtDNA of interest. In the next generation, hermaphrodites expressing GFP in germline and pharynx are selfed to create final vehicle strains, which are homozygous at GPR-1(OE) and pharyngeal GFP marker loci and contain the mtDNA of interest. Step 2-transferring mtDNAs into different nuclear backgrounds. Each of the wild strain males is mated to the vehicle strains. F_1_ progeny with distinct chimeric pharyngeal fluorescent pattern are singled from each cross and allowed to self to create a progeny with unique mitonuclear combination. **B. Pairwise divergence between the 6 wild isolates in the exchange panel used in high throughput phenotypic assay.** Bottom right nuclear genome divergence, top left mitochondrial genome divergence. Pairwise strain divergence is reported in unit of 10^−3^ per site.

During the second steps of the strain construction process, we observed the absence of chimeric progeny in certain crosses despite successful mating, i.e., the presence of heterozygous F_1_ males. We hypothesized that this lack of chimeric animals was caused by parental-effect toxicity loci, such as *peel-1/zeel-1* (Seidel, et al. 2011) and *pha-1/sup-35* (Ben-David, et al. 2017). We found that the failure of generating chimeric progeny correlated with variation at *pha-1/sup-35*, suggesting that the toxic N2 allele of *sup-35* locus was possibly fixed during vehicle strain generation. Such hermaphrodites produce and load maternal toxin SUP-35 into their oocytes. When crossed to males lacking the *pha-1/sup-35* locus, heterozygous offspring are ordinarily rescued by zygotic expression of the maternally transmitted *pha-1* allele, but chimeric offspring lack this maternal *pha-1* haplotype in the cell lineage carrying only the paternal genome. The result is embryonic arrest. To overcome this, we used RNAi to suppress *sup-35* expression and successfully constructed 323 of 324 possible combinations, including reintroduction of each mtDNA into its matched nuclear background as a control. One combination, the ECA2191 mitochondrial genome in the N2 nuclear background, resisted construction.

### Mitonuclear epistasis contributes to phenotypic variation in worm development

To assess the contribution of mitonuclear epistasis to phenotypic variation, we measured body size after 48 hours of growth from L1 arrest as a proxy for developmental rate in a subset of genotypes (6 nuclear × 6 mtDNA). We selected strains from six haplogroups to quickly assess how mitonuclear interactions affect phenotype, with the rationale that greater genetic divergence would likely yield more pronounced phenotypic differences. The sequence divergence between each pair of nuclear and mitochondrial genomes is shown in **Figure 2B**. Strains with the ECA2367 nuclear background grew slowly and failed to produce enough age-synchronized embryos within the assay timeframe (see **Methods**) and so were excluded from the final phenotypic dataset. In the end, we phenotyped 90 strains (30 genotypes × 3 independently derived mitochondrial exchange strains each) in a control condition with standard laboratory growth temperature (20°C), and under five different conditions hypothesized to tax mitochondrial function: three heavy-metal exposures (copper chloride, nickel chloride, and zinc sulfate) and one pesticide (paraquat) at 20°C, and an elevated temperature (25°C).

Independent of mitochondrial haplotypes (MT term), nuclear genotypes (N term) contribute to phenotypic variation across conditions (E term) (**Figure 3A**). For instance, among the strains with the same mtDNA, the ECA1229 nuclear background generally displayed a larger body size than the average, whereas ECA1493 exhibited a smaller body size than the average in most but not all conditions (**Figure 3A**). Departures from this pattern, such as the relatively small size of ECA1229 worms at 25°C, and relatively large size of ECA1493 worms in paraquat, illustrate the strong genotype-by-environment interactions present in the data (**Table 1**). The different mitochondrial haplotypes resulted in varied phenotypic effects across different nuclear backgrounds. Across nuclear backgrounds, each mitochondrial genotype sometimes increased worm length and sometimes decreased it, and no mitochondrial haplotype was associated with body size on average across nuclear backgrounds (**Figure 3A**). We observe a highly significant three-way MT × N × E interaction (P << 2.2×10^−16^) (**Table 1**), and highly significant MT × N epistasis within each of the six conditions (**Table 2**).

**Figure 3.**
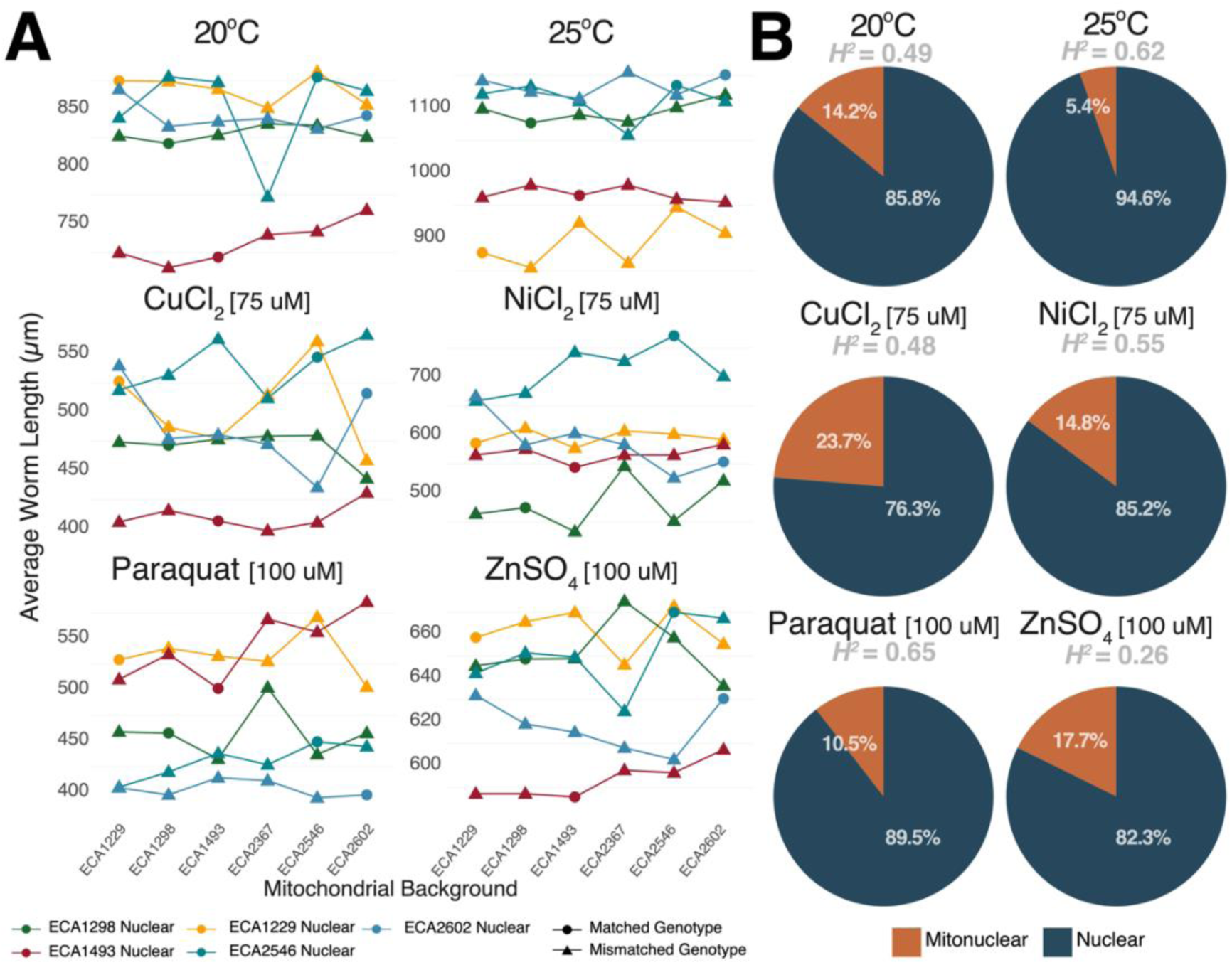
**A. Mitonuclear epistasis influences phenotypic variation**. Changes in phenotypes are presented as interaction plots, where each colored line follows the changes in body size of a single nuclear genetic background when paired with 6 different mt haplotypes, as indicated on the x-axis, under six different conditions. The ordering of mitotypes does not reflect genetic relatedness. Note that the scale of the y-axis is different for each media condition. **B. Mitonuclear epistasis contributes to genetic variance.** For each of the six conditions, we partitioned the heritable component of phenotypic variance into its constituents: main effect of mitochondrial genotype (not plotted because not detected), main effect of nuclear genotype (blue), and mitonuclear interactions (orange).

**Table 1.** Mitochondrial x nuclear x environment interactions (mt x n x e)

| Effect | d.f. | Models | P-values |
| --- | --- | --- | --- |
| Nuclear | 1 | Full, N | 0.896 |
| Mitochondrial | 1 | Full, MT | 1 |
| Condition | 5 | Full, E | $1.17 \times 10^{-14}$ |
| Nuclear × Condition | 1 | Full, N × E | $4.44 \times 10^{-55}$ |
| Mitochondrial × Condition | 1 | Full, MT × E | 1 |
| Nuclear × Mitochondrial | 1 | Full, N × MT | $6.01 \times 10^{-6}$ |
| Nuclear × Mitochondrial × Condition | 1 | Full, N × MT × E | $< 2.2 \times 10^{-6}$ |

**Table 2.** ANOVA tables showing nuclear, mitochondrial and mitonuclear effects on phenotype.

| Condition | EFFECT | Models | d.f. | P-values |
| --- | --- | --- | --- | --- |
| 20C | Nuclear | Full, N | 1 | $2.32 \times 10^{-9}$ |
|  | Mitochondrial | Full, MT | 1 | 1 |
| | mt × n interactions | Full, N × MT | 1 | $1.18 \times 10^{-144}$ |
| 25C | Nuclear | Full, N | 1 | $1.79 \times 10^{-14}$ |
|  | Mitochondrial | Full, MT | 1 | 1 |
| | mt × n interactions | Full, N × MT | 1 | $2.95 \times 10^{-102}$ |
| Copper | Nuclear | Full, N | 1 | $8.32 \times 10^{-7}$ |
|  | Mitochondrial | Full, MT | 1 | 1 |
| | mt × n interactions | Full, N × MT | 1 | $1.41 \times 10^{-261}$ |
| Nickel | Nuclear | Full, N | 1 | $4.05 \times 10^{-9}$ |
|  | Mitochondrial | Full, MT | 1 | 1 |
| | mt × n interactions | Full, N × MT | 1 | $3.64 \times 10^{-165}$ |
| Paraquat | Nuclear | Full, N | 1 | $3.83 \times 10^{-11}$ |
|  | Mitochondrial | Full, MT | 1 | 1 |
| | mt × n interactions | Full, N × MT | 1 | $2.03 \times 10^{-229}$ |
| Zinc | Nuclear | Full, N | 1 | $3.82 \times 10^{-8}$ |
|  | Mitochondrial | Full, MT | 1 | 1 |
| | mt × n interactions | Full, N × MT | 1 | $2.96 \times 10^{-69}$ |

Across environmental conditions, estimated broad-sense heritability (*H^2^*) ranged from 26% to 65% (**Figure 3B**). We further partitioned the contribution of each genetic term (nuclear genotype, mitochondrial genotype, and mitonuclear epistasis) to genetic variance (*V_G_*) within each growth environment (**Figure 3B**). Unsurprisingly, nuclear genotype explained the majority of variance in worm size. The main effect of mitochondrial genotype did not explain any of genetic variance, nor was it significant in any condition (**Table 2**). This observation aligns with results from yeast (Paliwal, et al. 2014; Nguyen, et al. 2020) and flies (Zhu, et al. 2014; Mossman, Ge, et al. 2019); the mitochondrial main effects were non-significant in all conditions, with mitochondrial effects instead dependent on nuclear background. Indeed, mitonuclear interactions explained a significant proportion of genetic variance, ranging from 5.4% to 23.7% within each environment. Overall, mitonuclear interactions are widespread across all conditions.

To understand how epistatic interactions were distributed among strains, we analyzed each pairwise mitochondrial exchange: each set of four strains consisting of two matched mitonuclear combinations and two mismatched mitonuclear combinations **(Figure 4A, Table S8).** In 58% of mitonuclear swaps (35 out of the 60 viable strain quartets), we observed a significant epistatic effect. Paraquat exhibited the highest rate, with 90% of mitochondrial exchanges showing significant epistatic effects, followed by high temperature (70%), aligning with the notion that high temperatures unveil mitonuclear interactions, underscoring the importance of mitochondrial function in thermal responses. Some wild isolates were particularly sensitive to changes in their mitonuclear combination. ECA2602 and ECA1229 were each involved in significant epistatic effects in every condition. The ECA2546-ECA1229 and ECA2546-ECA2602 pairs showed significance in 5 of 6 conditions, all but copper chloride (**Table S8**).

**Figure 4.**
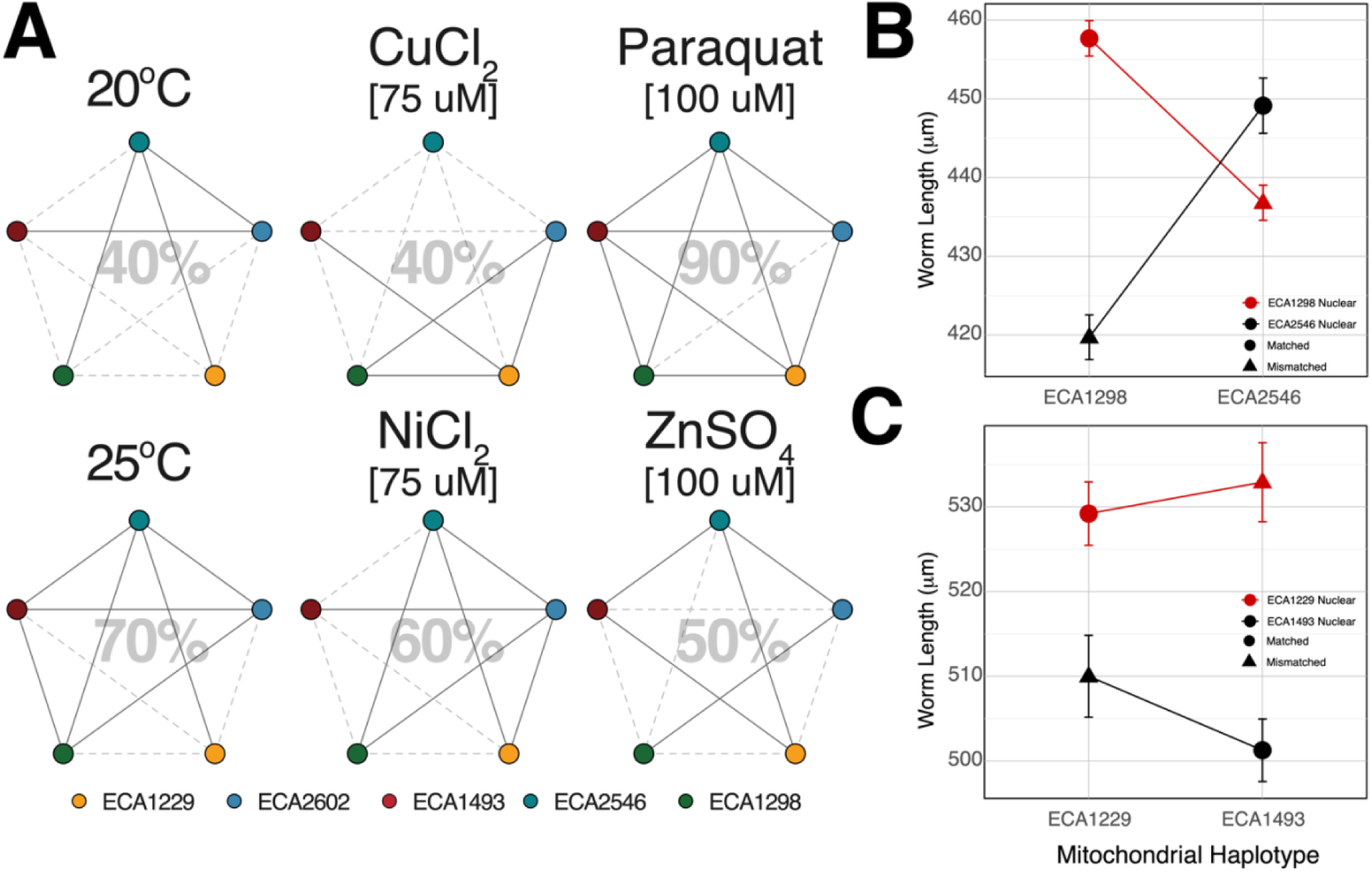
A. Frequency of mitonuclear epistasis. Bold lines connect the circles when significant mitonuclear epistasis was observed following an exchange of mtDNA between the two strains (two-way ANOVA, P < 0.05). mtDNA exchanges that did not reveal epistasis are shown as dotted gray lines. The percentage of all significant epistasis tests is shown for each condition. **B.** Representative case where matched genotypes grew faster than mismatched genotypes in Paraquat **C.** Representative case where matched genotypes grew smaller than mismatched genotypes in Paraquat. Error bars represented the standard error of the mean phenotypes.

Next, we examined if the phenotypes of matched mitonuclear combinations differed in a consistent direction from those of mismatched combinations; if matches confer robust developmental phenotypes, we expect matches to result in greater worm lengths in our assay. Among the strain pairs that showed significant epistatic interactions, we observed cases where matched nuclear genotypes with their own mitochondrial genomes resulted in longer worms compared to those with a foreign mitochondrial genome (**Figure 4B**, **Figure S3**). However, in many other cases, mismatched genotypes exhibited larger body sizes than their matched counterparts (**Figure 4C, Figure S3**). Given that the nuclear genome exerted the most significant influence on phenotype, we turned comparing the phenotypes of mismatched strains to matched combinations amongst strains that shared the same nuclear DNA (**Table S9**, **Figure S4)**. In our analysis, 14.7% (22 of 150) of matched strains were at least 5% larger than their mismatched counterparts, while mismatched strains were at least 5% larger in only 16 of 150 cases; in the majority of cases (74.7%, or 112 of 150), there was no statistical difference between mismatched genotypes from the matched genotypes or the genotypes were not at least 5% different in size from each other. While deciphering the pattern of coadapted mitonuclear combinations proves challenging and unpredictable due to the environment sensitivity of mitonuclear interactions, the nuclear background of ECA2546 was negatively affected by mtDNA swaps across all conditions (**Table S9, Figure S4**), offering promising avenues for further molecular studies.

## Discussion

In this study, we establish *C. elegans* as a powerful model for investigating mitonuclear interactions by leveraging the natural genetic diversity in both mitochondrial and nuclear genomes. Our panel of mitonuclear cybrids allows us to explore the interplay between mitochondrial and nuclear genomes and how these interactions influence phenotypic traits and responses to environmental challenges.

We found that *C. elegans* segregates more than 1,000 polymorphic sites within protein-coding regions of the mitochondrial genome. This level of variability is significantly higher than that reported in other model organism natural variation resources with mitochondrial variation data, including *Saccharomyces cerevisiae* (384 polymorphic sites) (De Chiara, et al. 2020) and in *Drosophila melanogaster* (231 sites) (Bevers, et al. 2019). The latter represents 169 lines of the Drosophila Genetic Reference Panel, all of which derived from a single collection site in North Carolina, USA (Mackay, et al. 2012), and the global mitochondrial genetic diversity of *D. melanogaster* may be higher than that of *C. elegans*. Regardless, *C. elegans* mtDNA is an outstanding model for investigation of how mitochondrial genotypes influence and interact with nuclear backgrounds.

The evolutionary pattern of *C. elegans* mtDNA appear to be marked by frequent recurrent mutations. Unlike outcrossing species, where sexual reproduction and recombination can lead to the introgression of beneficial nuclear alleles or even the complete purge of faulty mitochondrial genomes (Havird, Hall, et al. 2015), selfing species have limited opportunities for such genetic rescue to offset harmful mutations and may instead rely on accumulating compensatory mutations directly within mtDNA (Princepe and de Aguiar 2024). This could lead to an accelerated mutation-driven diversification, creating distinct evolutionary trajectories compared to outcrossing species. Future studies comparing mtDNA diversity between selfing and outcrossing species could help determine whether restricted nuclear gene flow in selfing lineages contributes to greater mtDNA diversity. The independent evolution of androdioecy three times within *Caenorhabditis* (Felix, et al. 2014) provides a valuable opportunity to directly compare mtDNA diversity in selfing versus gonochoristic species. One major challenge we encountered was polarizing mtDNA mutations in *C. elegans* with its closest available outgroup, *C. inopinata*, which diverged many tens of millions of generations ago (Kanzaki, et al. 2018). The high sequence divergence (dS = 6.72) made it impossible to confidently estimate the root of the *C. elegans* mitochondrial tree or the ancestral state of specific nucleotides or residues. *C. tropicalis* and *C. briggsae* have less divergent gonochoristic sister species (Woodruff, et al. 2010; Felix, et al. 2014), potentially allowing for direct comparisons without the deep divergence seen between *C. inopinata* and *C. elegans* in this study (Kanzaki, et al. 2018). This closer relationship enables better mutation polarization and more accurate reconstruction of ancestral states.

Perhaps the most intriguing finding is that HyPhy analysis corroborated our recurrent mutations results, indicating that the 94^th^ codon in *nduo-1* may have undergone adaptive changes; three different amino acid residues – phenylalanine, leucine, and isoleucine – all segregate at this site within *C. elegans*. To assess whether these recurrent mutations occur in conserved or variable regions, we aligned all available ND1 proteins in the *Caenorhabditis* genus from NCBI (accession numbers listed in **Table S10**) and examined the alignment (Figure **S5A**). Within the genus, residue 94^th^ is conserved as phenylalanine (F) in all species except *C. inopinata*, which carries a leucine (L) codon. Although this substitution preserves hydrophobicity, it alters the aromatic character that could be critical for electron transfer efficiency or protein-protein interactions. Expanding our analysis to include more distantly related nematodes reveals greater variability at residue 94 (**Figure S5B**). Using the ConSurf server (Ashkenazy, et al. 2010; Celniker, et al. 2013; Ashkenazy, et al. 2016; Yariv, et al. 2023) we determined that this residue is more variable among nematode ND1 peptides, with isoleucine (I), methionine (M), and valine (V) observed. This indicates that while the hydrophobic nature of the residue is maintained, the specific aromatic property of F is not universally required. Thus, within the *Caenorhabditis* genus, strong purifying selection appears to preserve residue 94, whereas across broader nematode lineages the site is semi-conserved and subject to modification. The L and I alleles in *C. elegans* occur near the tips of the mitochondrial phylogeny (**Figure S5C**), suggesting that these variants may confer transiently beneficial properties. Hypermutability remains an alternative explanation.

Complex I (NADH: ubiquinone oxidoreductase) has a fascinating structure, with ND1 playing a pivotal role at the junction of its two arms—a hydrophilic arm in the matrix (housing the electron transfer chain) and a membrane arm embedded in the inner mitochondrial membrane (containing all mitochondrially encoded subunits). This interesting placement makes ND1 essential for Complex I assembly and function as it directly contacts nuclear encoded mitochondrial proteins, such as NDUFS2, NDUFS 7 and NDUFS8 to support electron transfer and proton translocation (Hirst 2013; Parey, et al. 2020). To investigate whether the 94^th^ codon is located at an interacting interface, we aligned homologous protein sequences from species with well-characterized Complex I structures, including *Escherichia coli* (Kolata and Efremov 2021), *Thermus thermophilus* (Baradaran, et al. 2013), *Homo sapiens* (Stroud, et al. 2016), *Bos taurus* (Jones, et al. 2017), and yeast *Yarrowia lipolytica* (Parey, et al. 2021) (**Figure S6**). We inferred the transmembrane topology of NDUO1 based on a well-established topology model determined by protein fusion techniques (Roth and Hagerhall 2001) and utilized loop topology data from *E. coli* NuoH (Kervinen, et al. 2006). Based on this model, the 94^th^ residue of *C. elegans* NDUO1 is located in the loop connecting transmembrane helices 2 and 3 (TMH2-3^ND1^ loop), which is positioned in the intermembrane space (IMS) rather than the inner mitochondrial matrix (Roth and Hagerhall 2001), suggesting that residue 94 does not interact with matrix-facing nuclear-encoded subunits. However, its location in the IMS does not make it functionally irrelevant. The TMH2-3^ND1^ loop has been shown to play an important structural role in stabilizing TMH1^ND1^, another helix that interacts with the ND3 subunit in the membrane arm of Complex I (Padavannil, et al. 2021). Furthermore, residue 94 may facilitate interactions with assembly factors such as NDUFAF3, NDUFAF4, or TIMMDC1, which are known to assist in ND1 and ND3 integration into Complex I (Sanchez-Caballero, et al. 2016). It is plausible that variation at this site could have substantial structural and functional consequences, potentially impacting Complex I assembly, and biogenesis. Further functional studies would be necessary to determine whether 94^th^ residue in NDUO1 confer conditional advantages in energy metabolism or stress resistance in *C. elegans*.

In this study, we aimed to construct a comprehensive panel of mitonuclear cybrids in *Caenorhabditis elegans* to explore mitonuclear interactions. We employed RNA interference to suppress *sup-35* expression, a strategy that enabled the successful construction of 323 cybrid combinations by overcoming genetic incompatibilities. However, this approach did not succeed in all cases. One specific combination, between the nuclear genome of the N2 strain and the mitochondrial genome of ECA2191, resisted construction even with *sup-35* suppression. This persistent failure underlines the complexity of mitonuclear interactions and suggests the involvement of additional, unidentified factors governing nuclear-mitochondrial compatibility in *C. elegans*, perhaps sensitized by GPR-1 overexpression*. Medea* elements like *sup-35* (Ben-David et al. 2017) are pervasive in *Caenorhabditis* genomes (Ben-David, et al. 2021; Noble, et al. 2021) and could create lethal combinations in specific mitochondrial or maternal cytoplasmic backgrounds (Noble, et al. 2021; Pliota, et al. 2024).

To generate the mitonuclear strain panel, we employed the GPR-1 overexpression construct to exchange mitochondrial DNA between *C. elegans* strains. However, each vehicle strain, which we used to pass a mitochondrial genome into its final cybrid strain, also passes other cytoplasmic factors, including small RNAs with potential transgenerational phenotypic effects (Baugh and Day 2020). Such effects are potentially confounded with mitochondrial genotype in our experiment. Effects mediated by small RNAs in *C. elegans* typically fade after 3–5 generations (Rechavi, et al. 2011; Moore, et al. 2019) but in some cases persist for many generations (Vastenhouw, et al. 2006). Several factors reduce our concern about this issue. First, each strain was generated under identically controlled conditions without any of the external stressors known to trigger sRNA responses, such as parasites (Rechavi, et al. 2011), starvation (Rechavi, et al. 2014), or high temperature (Schott, et al. 2014). Second, maternal cytoplasmic effects should be confounded primarily with mitochondrial main effects, which we did not detect. Third, because we required only the mitochondrial genome from each vehicle strain, we allowed their nuclear genomes to be genetically heterogeneous, segregating N2 and focal-strain genotypes; consequently, each of the three independent replicates for each genotype (and each of the 18 that share a mitochondrial genotype) was derived from a genetically unique representative of the vehicle strain. To the extent that either maternal nuclear genotype or maternal environment influences the cytoplasmically transmitted small RNA pool, phenotypic effects of that pool would largely contribute to within-genotype variance rather than to the among-genotype variance that we detect as mitonuclear epistasis. Finally, the individual worms subjected to phenotyping had been propagated for five or more generations before assays. This propagation period likely helped “reset” any transient small RNA effects, reducing the chances of their persistence influencing our phenotypic measurements. While we cannot completely rule out the persistence of sRNAs in our strain collection, the stochastic nature of their inheritance, combined with our propagation strategy, makes it unlikely that they significantly affected the phenotypes we observed. Nonetheless, future studies should aim to disentangle mtDNA effects from any potential residual sRNA contribution to phenotypes.

Our study provides limited evidence for mitonuclear coadaptation in *C. elegans*, despite expectations that the selfing mating system would favor the evolution of coadapted mitonuclear interactions. While we hypothesized that strains with matched mitonuclear genomes would show greater resistance to stressors, such as heavy metal exposure, compared to mismatched cybrids, the results were mixed (**Figure 4B-C, Figure S4-5**). Matched combinations did not consistently perform better under stress conditions; some mismatched combinations displayed larger body sizes, indicating greater stress resistance, while others were smaller and more susceptible. These varied outcomes suggest that mitonuclear coadaptation may be highly context-dependent, with certain environmental conditions potentially favoring specific combinations. Interestingly, we observed the highest occurrence of matched genotypes conferring a growth advantage in copper chloride (**Figure S4**), with size differences of up to 15.6% —the largest effect size across all conditions. Copper exposure appeared to favor matched mitonuclear combinations in three nuclear backgrounds: ECA1229, ECA2546, and ECA2602. The underlying reason for this remains unclear, as we lack sufficient knowledge of the evolutionary histories and ecological contexts of these strains to draw definitive conclusions.

While these findings highlight the impact of mitonuclear interactions on phenotypes, they also reveal the limitations of using simple traits, like worm length, as proxies for fitness. Because mitonuclear interactions are sensitive to environmental conditions, meaningful selection for optimal interactions likely occurs in ecologically relevant settings (Nguyen, et al. 2020), and phenotypic outcomes may vary depending on the specific environment. Future research should focus on more comprehensive life-history traits, such as fecundity, time to sexual maturity, and longevity, along with molecular phenotypes like ATP production and ROS levels. Combining these with transcriptomic and proteomic analyses could reveal the pathways influenced by specific mitonuclear combinations and provide deeper insights into the functional consequences of these interactions.

In summary, this study demonstrates the substantial polymorphism of *C. elegans* mitochondrial DNA and its influence on mitonuclear interactions and phenotypes. Our findings highlight the importance of accounting for both genetic and environmental contexts when examining the evolutionary dynamics of mitonuclear epistasis. *C. elegans* is a valuable model for future research focused on unraveling the complexities of mitonuclear coevolution and its role in local adaptation.

## Materials and Methods

### Mitochondrial molecular evolution

The soft-filtered VCF file for all 1,385 strains (20220216 release) was obtained from the *Caenorhabditis* Natural Diversity Resource (Crombie, et al. 2024). Each strain has The VCF file was first filtered to retain only high-quality variants marked with “PASS” flag (QUAL >30, QD >20, SOR <5, FS<100, DP >5) using bcftools (version 1.14) (Danecek, et al. 2021). At the sample-level, we retained 1,372 strains with <10% mitochondrial heterozygosity using vcftools (version 0.1.16) (Danecek, et al. 2011). Because the VCF was generated using a diploid variant calling model, we observed 45 heterozygous genotype calls across 27 strains at 40 unique mitochondrial sites. These sites were included in downstream analyses, but heterozygous calls were converted to missing data. While some of these may result from sequencing error, they may also reflect true heteroplasmic states of the mitochondrial genome. Each strain has at least 14x nuclear genome coverage, the median mitochondrial genome coverage is 597x, with a median site-specific variant quality score of 209,812. To ensure reliable variant calls, we set a quality score threshold of 1,000 for SNPs and a more stringent threshold of 10,000 for indels, accounting for their higher susceptibility to genotype calling errors, which matched our assessments from manual inspection od read alignments. To generate the final VCF file, we removed one multiallelic intergenic indel spanning multiple bases, as it interfered with downstream sequence alignment. Finally, SNPEff (version 5.1) (Cingolani, et al. 2012) was used to annotate and predict the effects of single nucleotide polymorphisms using the invertebrate mitochondrial genetic code. While we observed many codons with triallelic sites, only 12 codons showed variation at multiple nucleotide positions (**Table S11**). In such cases, SnpEff may yield inaccurate annotations, as it evaluates each variant independently relative to the reference sequence and does not account for phased combinations. We identified and corrected one clear misannotation at mtDNA position 3036, which should have been classified as synonymous. Changes in most of the other codons are unambiguously synonymous or nonsynonymous, and all are included in **Table S11**.

Consensus sequences were generated for each sample based on Reference genome FASTA (WS283) using bcftools (version 1.14) (Danecek, et al. 2021). All processed FASTA files were concatenated into a single file and aligned using MAFFT (version 7.475) (Katoh and Standley 2013). Identical haplotypes were identified using the IdenticalHaplotype.R script (https://github.com/Tuc-Nguyen/mitonuclear-elegans).

To investigate the nuclear population structure of all 540 isotypes, we performed a PCA on our VCF file using PLINK (v1.9) (Chang, et al. 2015). The nucleotide divergence for both mitochondrial and nuclear genomes between strain pairs in **Figure 2** were calculated using VCFtools (version 0.1.16) (Danecek, et al. 2011). A sliding window approach with a 1,000 bp window size was used to count the number of variants in each window, and the average pairwise nucleotide differences across all windows were estimated as pairwise divergences. It is important to note that nuclear divergences are likely underestimated due to read alignment challenges in hyperdivergent regions (Lee, et al. 2021).

#### Recombination Analysis

To examine potential mitochondrial recombination, we analyzed the final whole mitochondrial genome alignment using various methods available in RDP5 (Recombination Detection Program) (Martin, et al. 2021). Specifically, we applied six different detection methods: 3Seq (Lam, et al. 2018), GENECONV(Padidam, et al. 1999), MaxChi (Smith 1992), Chimaera (Posada and Crandall 2001), SiScan (Gibbs, et al. 2000), and the original RDP method (Martin and Rybicki 2000) with the default settings on circular genomes.

*Phylogenetic inference* among *C. elegans* isolates was performed on the alignment file containing 367 unique sequences using IQ-TREE (version 1.6.12) (Nguyen, et al. 2015). The best-fit model TN+F+R3 (Tamura-Nei with empirical base frequencies and FreeRate model with 3 categories) was selected using ModelFinder (Kalyaanamoorthy, et al. 2017) based on the lowest BIC score. This model accounts for transition-transversion biases, unequal base frequencies, and heterogeneous site rates. The final model parameters included empirical nucleotide frequencies (A: 0.314, C: 0.089, G: 0.149, T: 0.448) and substitution rates (A↔C: 1.000, A↔G: 9.29239, A↔T: 1.000, C↔G: 1.000, C↔T: 16.3349, G↔T: 1.000). Rate heterogeneity was modeled with three FreeRate categories (R3), with site proportions and relative rates as follows: Category 1 (88.14%, 0.2283), Category 2 (11.31%, 5.449), and Category 3 (0.5529%, 32.99). Additional details on model parameters, rate heterogeneity, and tree statistics can be found in **Supplemental File 1.** The best-scoring ML tree had a final log-likelihood of −36,048.0554 after 111 iterations and was visualized using the ggtree package (version 3.1.3.993) in R version 4.0.2 (Yu, et al. 2017). We used the HaploDiagnosticMutations.R script (https://github.com/Tuc-Nguyen/mitonuclear-elegans) to infer the number of mutational changes per branch across the mitochondrial phylogeny. In brief, we mapped the 472 putatively functional variants onto the tree using stochastic character mapping via the *make.simmap* function from the *phytools* package (Revell 2024). This mapping produced detailed output that described the duration each branch spent in different states, i.e. branches exhibiting two mapping segments indicated a state change, providing the basis for our parsimony-like reconstruction. To find the minimum number of changes required by the tree, we then remapped all SNPs using a modified rate matrix (0.001×fixedq) and recorded the number of transitions per branch, allowing us to assign mutation events across the mitochondrial phylogeny.

#### Inferring selection pressure

We extracted gene sequences from each of the 367 unique mitochondrial genomes using the GenBank annotation of the *C. elegans* reference sequence (WS283). Sequences for each gene were specifically aligned using MAFFT (version 7.475) to prepare for subsequent analyses. The phylogenetic tree inferred from whole mitochondrial genome alignments was used to preserve relative evolutionary distances between haplotypes. We employed the HyPhy (version 2.5.64) software suite (Kosakovsky Pond, et al. 2020) to detect selection on all mitochondrial protein-coding genes. For all 12 coding genes, we employed the **M**ixed **E**ffects **M**odel of **E**volution (**MEME**) method to identify episodic positive selection at individual sites (Murrell, et al. 2012). Additional details on each analysis can be found in **Supplemental File 2**.

#### Polarizing C. elegans mutations with Caenorhabditis inopinata

To infer evolutionary relationships with the sister species, we first assembled *C. inopinata* mitochondrial genome from both short reads and PacBio reads (Kanzaki, et al. 2018) obtained from GenBank (Accession: DRZ086531), as the *C. inopinata* reference genome lacked mitochondrial assembly data. An initial alignment was performed by mapping all raw PacBio reads to the reference genome (PRJDB5687, WS289 version) using minimap2 (version 2.22). Unmapped reads, presumed to contain mitochondrial genome reads, were extracted using samtools (version 1.14). These unmapped reads were subsequently aligned to the *C. elegans* mtDNA (WS283) to identify orthologous reads with minimap2. The *C. inopinata* mtDNA was *de novo* assembled from these orthologous reads with Flye (version 2.9.2), producing a single contig of 27,124bp, seemingly a fusion of two 13kb contigs due to the circular nature of mtDNA. Pairwise sequence alignment between *C. elegans* mtDNA and the contig was performed using BLAST+ (version 2.13.0), identifying an aligned hit 13,820bp long with 84% match. The long-read assembled contig of 13,883bp was generated by extracting the aligned 13,820bp sequence along with the subsequent 63bp preceding the sequence repetition. PacBio and short reads were mapped back to the 13,883bp contig using minimap2 (version 2.22) and bwa (0.7.17), respectively. Aligned reads were subsequently *de novo* assembled with SPAdes (version 3.15.0), generating a 14,276 contig. Using MUMmer (version 3.23), repetitive AT-rich regions were identified at the beginning and end of the 14 kb contig. The 14 kb contig was then aligned back to the original 13,883 bp contig using MUMmer (version 3.23), resulting in a nearly identical (99.98%) alignment spanning 13,794 bases. Given that the length of *C. elegans* mtDNA is 13,794 bp, the final sequence for *C. inopinata* for downstream analyses was derived from this aligned region. Pairwise mitochondrial sequence divergences among *C. elegans* isolates and between *C. elegans* (N2) and *C. inopinata* were calculated using KaKs_Calculator version 3.0 (Zhang 2022). We *de novo* annotated the *C. inopinata* mitochondrial genome using MITOS2 (Al Arab, et al. 2017; Donath, et al. 2019) to obtain mitochondrial protein-coding gene sequences.

For each of the 367 unique *C. elegans* mitochondrial haplotypes, we extracted gene sequences based on the GenBank annotation of the reference sequence (WS283), translated them into amino acid sequences, concatenated them into a final protein-coding sequences, and aligned them with *C. inopinata*’s concatenated amino acid sequence using MAFFT (version 7.475). Once aligned, a maximum likelihood phylogenetic tree was generated using IQ-TREE, (version 1.6.12). Model selection with ModelFinder identified mtMet+F+R3 (Mitochondrial Metazoan substitution model with empirical state frequencies and FreeRate model with 3 categories) as the best-fit model based on Bayesian Information Criterion (BIC). Maximum-likelihood inference was performed with 1000 ultrafast bootstrap replicates, where a constrained phylogenetic tree was enforced to maintain the intra-*C. elegans* phylogeny while allowing the position of the *C. inopinata* branch to vary. Additional details on model parameters, rate heterogeneity, and tree statistics can be found in **Supplemental File 3**. The tree was visualized using the ggtree package (version 3.1.3.993) in R.

### Strain construction and maintenance

All strains used in this study were maintained in the lab at 20°C on modified nematode growth medium (NGMA) plates (1% agar and 0.7% agarose) seeded with OP50-1 *Escherichia coli*. The specific growth conditions for the high-throughput assay are described in the following section.

#### Mitonuclear cybrid worm construction

The 18 wild isolates in this study (ECA1129, ECA1229, ECA1298, ECA1493, ECA2041, ECA2151, ECA2191, ECA2367, ECA2546, ECA2602, ECA2607, ECA2609, ECA396, N2, NIC1799, MY1, QX1211, XZ1516) were purchased from the *Caenorhabditis* Natural Diversity Resource and the toolkit strain PD2218 from *Caenorhabditis* Genetic Center. Males of each of the 18 wild isolates were mated to hermaphrodites of the GPR-1(OE) toolkit strain PD2218, which carries two transgenes integrated at different positions on chromosome III: one drives expression of GFP-tagged GPR-1 in the germline (*ccTi1594[mex-5p::gfp::gpr-1::smu-1 3′ UTR, cbr-unc-119(+)],* III:680195) and the other drives GFP in the pharynx (*umnIs7[myo-2p::gfp + NeoR]*, III:9421936) (**Figure 2A**). F_1_ males were then crossed to the corresponding wild isolate hermaphrodite harboring the mtDNA of interest. In the next generation, hermaphrodites expressing GFP in germlines and pharynx were identified and allowed to self to create strains that are homozygous for the GPR-1(OE) and pharyngeal GFP loci and contain the mtDNA of interest. The nuclear genomes of these “vehicle” strains are heterogeneous mosaics of wild isolate and PD2218 genomes. Next, males of each of the 18 wild isolates were mated to each of the vehicle strains (**Figure 2A**). Progeny with desired GPR-1-induced chimerism, indicated by mosaic GFP expression in the pharynx, were singled and allowed to self to create a unique line. To facilitate crosses between lineages that differ in the presence/absence of *sup-35/pha-1* maternal-effect gene-drive locus (Ben-David, et al. 2017), worms were cultured on NGMA-RNAi plates (NGMA +1mM IPTG and 25µg/ml carbenicillin) seeded with bacterial lawn containing sjj_Y48A6C.3 (Kamath, et al. 2003), an HT115 *E. coli* clone that produces *sup-35* dsRNA, inducing RNAi suppression of the *sup-35* toxin. Each of the 36 mitonuclear combinations was generated three times independently, yielding 108 strains for phenotyping. Due to slow growth, the 18 strains with ECA2367 nuclear background were subsequently excluded from phenotyping assays.

### Phenotyping Assay

In each of three independent experimental blocks, 30 mitonuclear cybrid genotypes were phenotyped in a high-throughput assay using the Molecular Devices ImageXpress Nano microscope (Molecular Devices, San Jose, CA). As described in (Widmayer, et al. 2022), strains were cultivated for three generations on NGMA plates at 20°C in uncrowded conditions. Gravid hermaphrodite populations from the fourth generation were synchronously bleached. In each of the three assay blocks, the strains were randomly distributed into two distinct groups for bleaching. The composition of strains within the bleaching groups were different across the four experimental blocks. Following bleach synchronization, 50 embryos suspended in 50µl of K medium were aliquoted into 96-well microtiter plates. Strain allocation to columns within the 96-well microplates was randomized, with each plate accommodating 10 strains, and each strain occupying 8 wells. Plates were sealed with gas-permeable sealing film (Axygen BF-400, Fisher Cat #: 14-222-043) and incubated in humidity chambers at 20°C with shaking at 170 rpm. After 24 hours, populations of developmentally arrested first larval-stage animals (L1s) were fed with of OD_600_100 HB101 *E. coli* bacterial culture along with pre-determined concentrations of heavy metals. The HB101 culture stock was prepared using the methods outlined (Widmayer, et al. 2022) under “Nematode food preparation.” The concentrations of heavy metals used in this study were derived from the dose-response curves established for copper (II) chloride, nickel chloride, zinc chloride, and paraquat as detailed in (Widmayer, et al. 2022). These concentrations were determined based on worm lengths falling within a range 100-300µm less than those in the control treatment. We added 25µl of the HB101 food and toxicant mix to the corresponding 96-well plates to simultaneously feed the arrested L1s at a final concentration of OD_600_10 and subject them to stressor exposure. All plates were then sealed, and placed in humidity chambers, and incubated for 48 hours at 20°C (or 25° where noted) with constant shaking. After 48 hours, a 10-minute treatment with sodium azide (50mM in 1X M9 Buffer) was applied to immobilize and straighten the worms immediately before imaging. The images were acquired using the Molecular Devices ImageXpress Nano microscope with a 2X objective. We followed the Methods described in (Widmayer, et al. 2022) to extract and filter animal measurements from images using CellProfiler software (Wahlby, et al. 2012), and *R/easyXpress* (Nyaanga, et al. 2021). These measurements comprised our raw phenotypic dataset.

### Quantitative genetics

Statistical analyses were performed in R version 4.2.1 (R Core Team 2022). The significance of nuclear and mitochondrial genetic components, environments, and their interactions were tested using Mixed-Effect models implemented in the lme4 with the restricted Maximum Likelihood (REML) estimation (Bates, et al. 2015). To determine the significance of each component and their interactions, we compared the fit of a full model that included all terms N + MT + E + (N × E) + (MT × E) + (MT × N) + (MT × N × E) to a reduced model where the term being evaluated was omitted. Likelihood Ratio Tests were used for model comparisons, where a significant chi-squared statistic indicates that the inclusion of the tested term significantly improves the model fit. Fixed effects were estimated for treatment (environment) and its interactions with nuclear and mitochondrial genotypes, while batch effects were incorporated as random effects to account for both global and within-environment variability. Since each experimental batch contained the same genotypes and each strain was phenotyped only once, strain effects are (by design) completely confounded with batch effects; consequently, a separate random effect for strain was not included due to model overparameterization. Similarly, within each environment we fitted random-effects models that included random effect batch term to control for batch-to-batch variation while testing the significance of nuclear, mitochondrial, and mitonuclear interactions. Model comparisons via LRTs provided the statistical evidence for the contribution of each component to variation in worm length.

#### Variance Component Analysis

The contributions of each genetic term to phenotypic variance within each environment were estimated from the full model: N + MT + (MT × N) + ε. The proportion of variance that can be explained by all genetic terms represents broad-sense heritability (*H^2^*). The proportion of each genetic terms relative to the total *V_G_* is reported in Figure 3. We also fitted random-effects models that included (1|Batch) to control for batch-to-batch variation while partitioning the variance due to nuclear, mitochondrial, and mitonuclear interactions.

#### Epistasis Frequency Analysis

To estimate the frequency of mitonuclear epistasis when exchanging mtDNA strains from the same or two different subpopulations, two-way ANOVAs were used to test the significance of mitonuclear interactions among each mtDNA exchange such that each test included 4 genotypes: two matched mitonuclear genotypes and two mismatched genotypes derived from the exchange of mtDNAs between the parental strains.

To assess the impact of disrupting naturally occurring mitonuclear combinations, we compared body size differences between each nuclear genotype’s matched strain and its five mismatched derivatives for all conditions. Specifically, we performed ANOVAs comparing the full mixed-effects model (including the mitochondrial genotype as a fixed effect) with a reduced model excluding the mitochondrial term, with the batch factor included as a random effect to account for batch variation.

## Author Contributions

T.H.M.N and M.V.R designed the project. THMN constructed the mitonuclear collection and collected high throughput data; DAK, ERW and OSW collected high throughput data, THMN and MVR performed statistical analyses; THMN wrote the manuscript. MVR revised and edited the manuscript.

## Data and Code Availability

Strains are available upon request from the corresponding authors. R and Bash scripts used in this study, along with all associated data, are available at https://github.com/Tuc-Nguyen/mitonuclear-elegans.

## Acknowledgements

We thank Claire Curtin and Kahlan Wilson for their help with strain husbandry and cryopreservation; Zoey Chen, Chenyu Wang, and Derin Çağlar for their technical assistance with the high-throughput assay; the David Fitch lab at NYU for providing RNAi reagents; and Timothy A. Crombie, Samuel J. Widmayer, and Erik Andersen for supplying wild *C. elegans* isolates and detailed protocols for bacterial food preparation and high-throughput assays. We are also grateful for the support from the NYU High Performance Computing Cluster, CaeNDR (supported by NSF CSBR 1930382), and the *Caenorhabditis* Genetics Center (supported by NIH OD010440). Lastly, we thank the members of the Rockman lab and Xiaoxue (Snow) Zhou for their valuable feedback and discussions.

---

**Figure S1.**
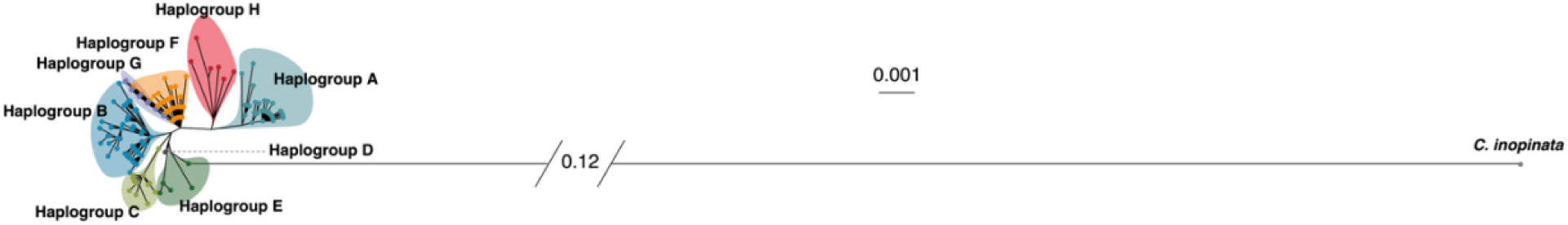
*Caenorhabditis inopinata* mitochondrial genome is substantially divergent from *Caenorhabditis elegans* mitochondrial genome. **Maximum Likelihood Phylogenetic tree** based on alignment of concatenated amino acid sequences from *C. inopinata* and 367 *C. elegans* isotypes.

**Figure S2.**
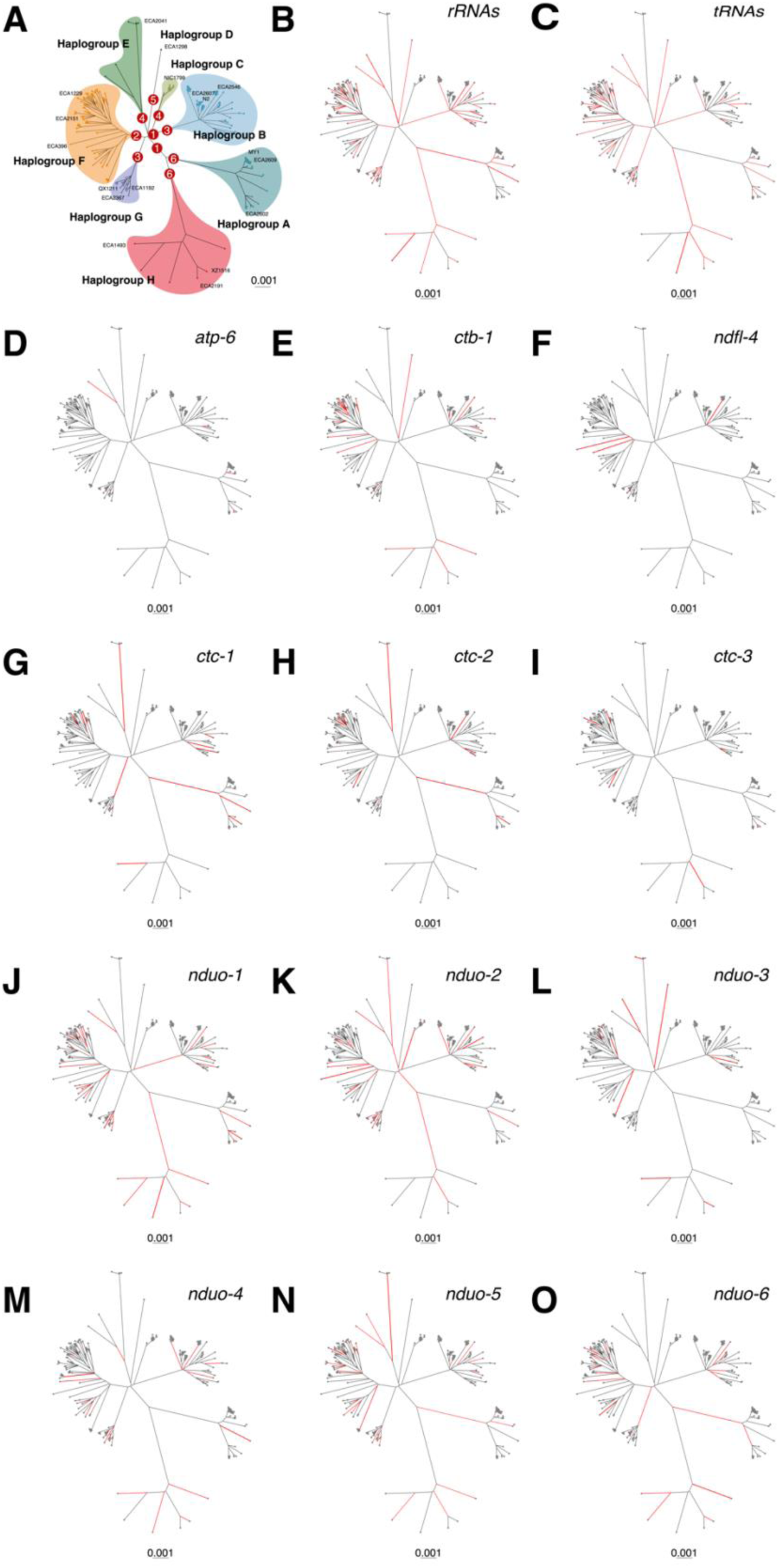
Phylogenetic distribution of mitochondrial functional variants. **A.** Number of functional changes (missense, rRNA, and tRNA) on the internal branches that connect mitochondrial haplogroups. **B-O Functional changes across mitochondrial protein-coding genes**. Branches exhibiting state transitions (mutations) are highlighted in red, while those with no detected changes remain in grey. The thickness of each branch is scaled to reflect the number of mutations, with thicker branches indicating a higher number of mutations.

**Figure S3.**
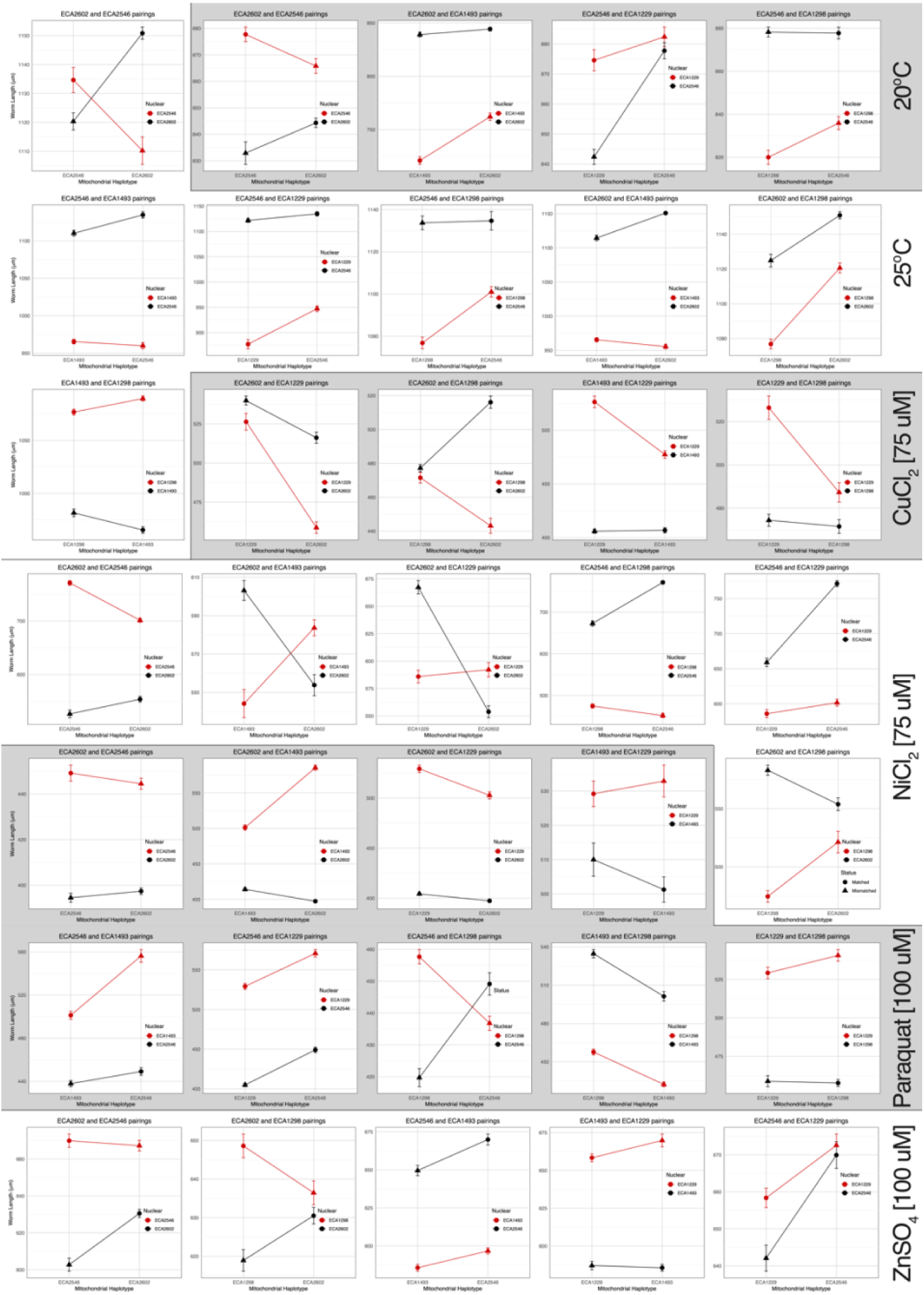
Interaction plots for strain pairs showing significant mitonuclear epistasis induced by mitochondrial swaps in all conditions. Each plot includes two matched mitonuclear combinations and two mismatched mitonuclear combinations. The shown quartets only represent cases where significant mitonuclear epistasis was detected (ANOVA, P < 0.05).

**Figure S4.**
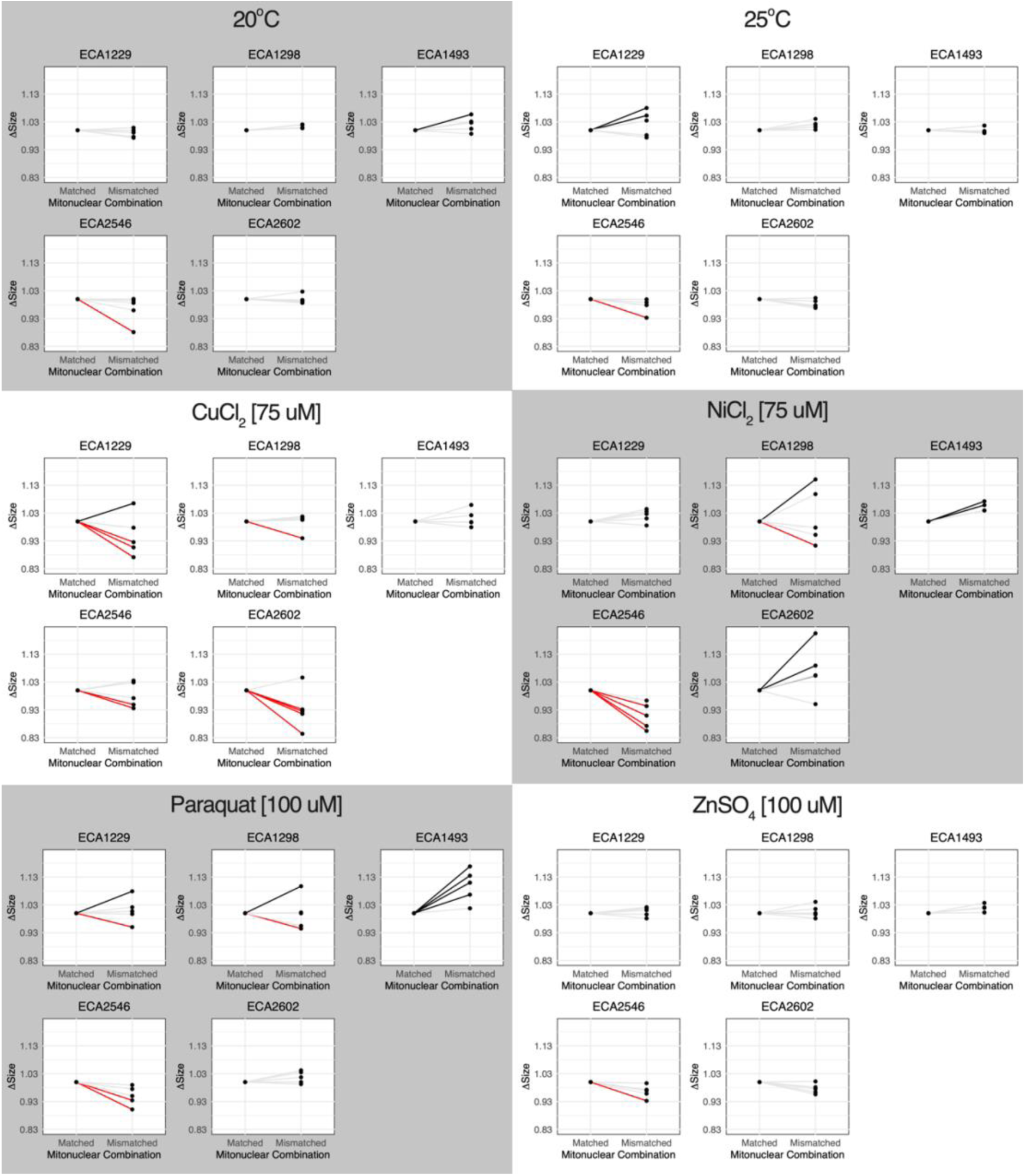
Relative phenotypic changes strains with mismatched mitonuclear genotypes to matched mitonuclear genotypes within the same nuclear background. Lines connecting the matched (reconstituted parental combination) and synthetic mitonuclear are colored to indicate a statistically significant reduction (red), increase (black), or less than 5% change (gray). See **Table S9** for statistics.

**Figure S5.**
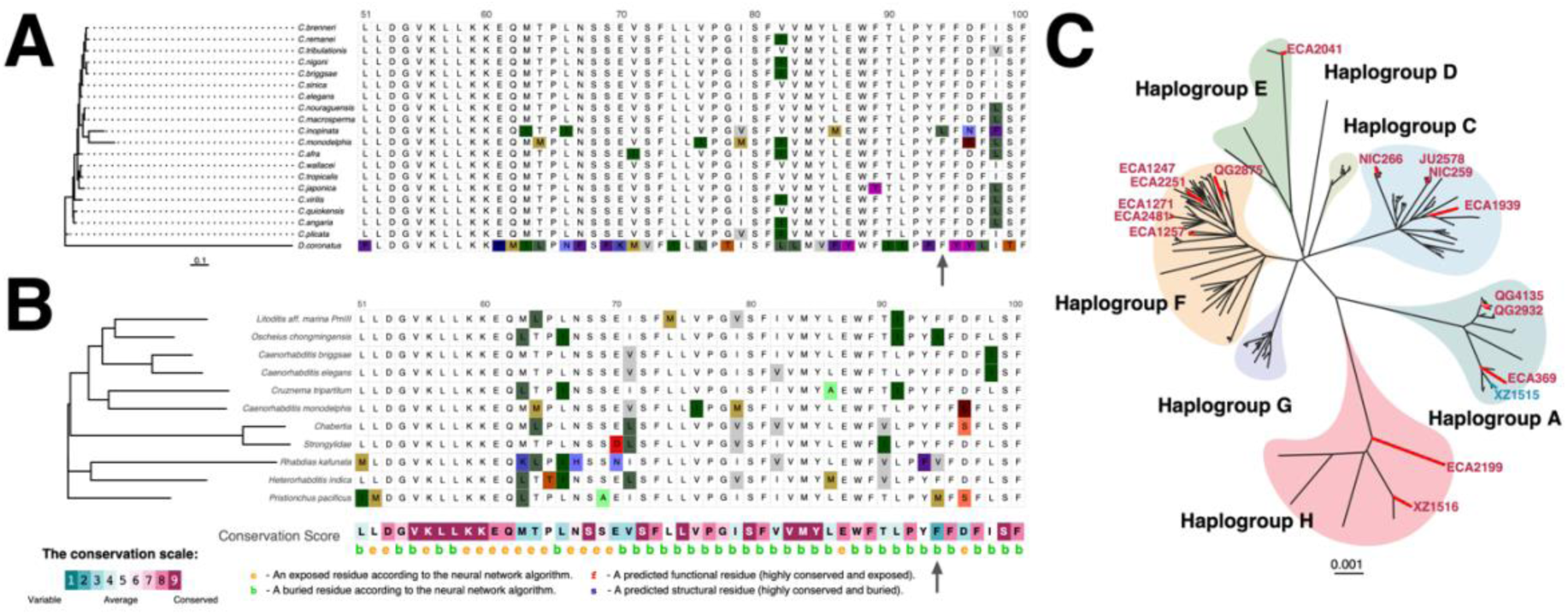
**A. The 94^th^ residue is highly conserved within *Caenorhabditis* genus.** Shown are amino acid residues from positions 50 to 100, with the 94^th^ residue indicated by a black arrow. Phylogenetic tree was generated based on whole gene alignment using mafft online server (Katoh, et al. 2017). Gene tree does not recapitulate species tree. **B. Amino acid conservation analysis of the mitochondrial peptide nduo-1 across a broader evolutionary scale.** Shown are amino acid residues from positions 50 to 100, with the 94^th^ residue indicated by a black arrow. Conservation analysis was performed using ConSurf server, which identified homologous sequences from UNIREF90 using the HMMER search algorithm. Pairwise identity thresholds were set to a maximum of 95% and a minimum of 80%. A multiple sequence alignment was built using MAFFT, and 12 representative sequences were selected to the final analysis. Phylogeny does not reflect species phylogeny. Conservation scores range from 1 (highly variable, teal) to 9 (highly conserved, purple) (Yariv, et al. 2023). **C. Distribution of the most recurrent mutation (mtDNA:2042) on the mitochondrial phylogeny.** Strains carrying alternative alleles are mapped onto the tree, with red indicating strains carrying the first alternative allele A (94F>94I) and blue representing strain with the second alternative allele C (94F>94L).

**Figure S6.**
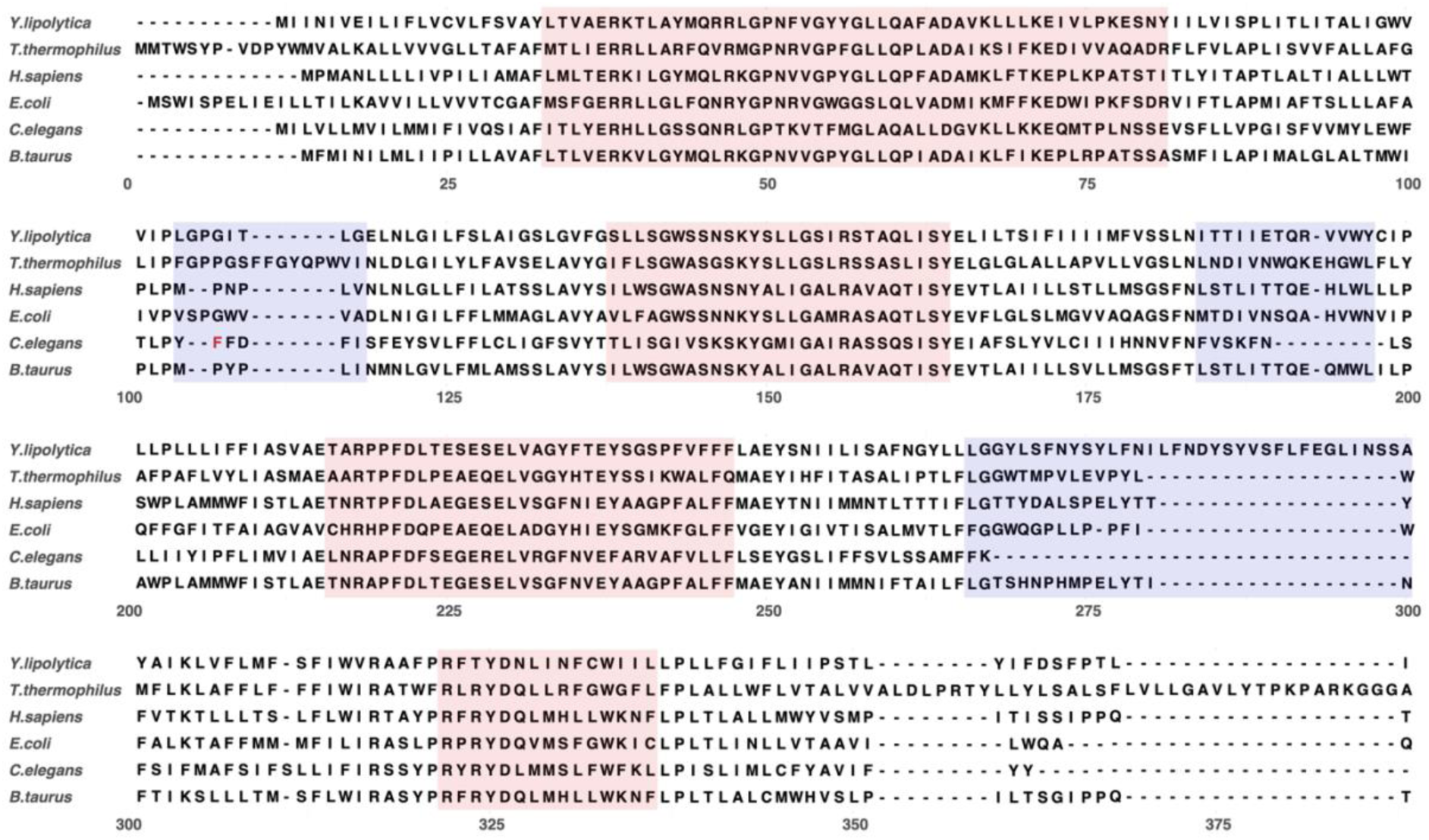
Sequence alignment of the complex I subunit ND1/NDUO1/NuoH/Nqo8. Residues belonging to the inner mitochondrial matrix or the inside ends of the predicted transmembrane helices are highlighted in red. Residues belonging to the intermembrane space or the outside of the predicted transmembrane helices are highlighted in purple. The 94^th^ codon of *C. elegans* NDUO1 is depicted in red. The alignment was carried out using mafft with default parameters. Accession numbers for sequences are (in order of appearance): CAC28089.2, AAA97945.1, CAA24026.1, CAA48367.1, AAO16387.1, CAA23997.1.

**Table S1. Strain information and mitochondrial haplotype classification.** Table includes strain names, isotypes, geographic coordinates (latitude and longitude), locations of isolation, mitochondrial haplogroups, and shared common haplotypes.

**Table S2. Intraspecific variant annotations of all mitochondrial SNPs segregating among *C. elegans* isotype reference strains**. Table includes the 1,458 mitochondrial sites that segregate among 367 C. *elegans* isotypes along with predicted amino acid changes using SNPEff (version 5.1). There are multiple rows for some sites because they segregate more than two variants.

**Table S3. Haplotype pairs differing by a single mitochondrial mutation among 367 isotypes.** Table highlights 144 single mitochondrial mutations that can be tested individually, including 63 missense mutations and 26 RNA mutations, to reveal the effects of specific mitochondrial changes. Highlighted cells reflect instances where there are single missense differences with the N2 reference genome.

**Table S4. Intra-isotype variants.** 28 mitochondrial sites that segregate amongst 832 other natural isolates that were not included in 540 isotypes.

**Table S5. Characterization of recurrent mutations on the mitochondrial phylogeny.** 86 SNPs (from the set of 472 missense or RNA-altering SNPs) that required at least 2 mutations on the tree are reported, along with the genes in which they are found, variant types, and minimum number of changes at that site required by the tree.

**Table S6. Diagnostic functional mitochondrial variants for each mitochondrial haplogroup. Recurrent mutations are highlighted.**

**Table S7. dN/dS ratio estimated by HyPhy for the internal branches and the tip branches for each mitochondrial coding gene.**

**Table S8. Strain pairs showing significant mitonuclear epistasis induced by mitochondrial swaps in all conditions**. For each of the strain pair, ANOVAs comparing the full mixed effect model with a model lacking the mitonuclear (N × MT) term were evaluated. The mitonuclear term was treated as a fixed effect, while batch factor was treated as a random effect.

**Table S9. The effect of mitochondrial DNA swap in each nuclear background across all environments.** To determine whether the difference among strains with matched and mismatched combinations is statistically significant, ANOVAs comparing the full mixed effect model with a model lacking the Mito term were evaluated. The Mito term was treated as a fixed effect, while batch factor was treated as a random effect.

**Table S10. Accession number for ND1 amino acid sequences from *Caenorhabditis* species and the outgroup *Diploscapter coronatus*.**

**Table S11. Codons that vary at multiple positions.** The table includes codon genotypes, their corresponding amino acids (based on the invertebrate mitochondrial genetic code), and the number of strains carrying each genotype. Red numbers indicate nucleotide positions with variants. The highlighted row denotes a case of SnpEff misannotation, where the change was predicted as a missense mutation, though the corresponding genotype is not observed in the population.

**Supplemental File 1**. Additional details on model parameters, rate heterogeneity, and tree statistics on intra *C. elegans* mitochondrial phylogeny

**Supplemental Files 2.** Additional details on model parameters, and statistics in all protein coding genes using Mixed Effect Model of Evolution in HyPhy.

**Supplemental File 3.** Additional details on model parameters, rate heterogeneity, and tree statistics on mitochondrially encoded amino acid phylogeny with *C. inopinata* outgroup.

