## Supplemental File 3 for "Abundant Recurrent Mitochondrial Mutations and Widespread Mitonuclear Epistasis in *Caenorhabditis elegans*"

IQ-TREE multicore version 1.6.12 for Linux 64-bit built Aug 15 2019
Developed by Bui Quang Minh, Nguyen Lam Tung, Olga Chernomor,
Heiko Schmidt, Dominik Schrempf, Michael Woodhams.

Host:    cm007.hpc.nyu.edu (AVX512, FMA3, 377 GB RAM)
Command: iqtree -s FINAL.fasta -m MFP -g constraint -bb 1000 -nt AUTO
Seed:    940353 (Using SPRNG - Scalable Parallel Random Number Generator)
Time:    Wed Mar 19 01:21:49 2025
Kernel:  AVX+FMA - auto-detect threads (48 CPU cores detected)

Reading alignment file FINAL.fasta ... Fasta format detected
Alignment most likely contains protein sequences
Alignment has 368 sequences with 3436 columns, 413 distinct patterns
119 parsimony-informative, 481 singleton sites, 2836 constant sites
           Gap/Ambiguity  Composition  p-value
   1  inopinata    0.26%    passed     98.99%
   2  AB1          0.26%    passed    100.00%
   3  BRC20067     0.26%    passed    100.00%
   4  BRC20263     0.26%    passed    100.00%
   5  CB4852       0.26%    passed    100.00%
   6  CB4856       0.26%    passed    100.00%
   7  CX11254      0.26%    passed    100.00%
   8  CX11264      0.26%    passed    100.00%
   9  CX11271      0.26%    passed    100.00%
  10  CX11292      0.26%    passed    100.00%
  11  CX11307      0.26%    passed    100.00%
  12  CX11314      0.26%    passed    100.00%
  13  DL226        0.26%    passed    100.00%
  14  DL238        0.26%    passed    100.00%
  15  ECA1069      0.26%    passed    100.00%
  16  ECA1185      0.26%    passed    100.00%
  17  ECA1187      0.26%    passed    100.00%
  18  ECA1191      0.26%    passed    100.00%
  19  ECA1192      0.26%    passed    100.00%
  20  ECA1194      0.26%    passed    100.00%
  21  ECA1200      0.26%    passed    100.00%
  22  ECA1202      0.26%    passed    100.00%
  23  ECA1206      0.26%    passed    100.00%
  24  ECA1208      0.26%    passed    100.00%
  25  ECA1212      0.26%    passed    100.00%
  26  ECA1216      0.26%    passed    100.00%
  27  ECA1223      0.26%    passed    100.00%
  28  ECA1225      0.26%    passed    100.00%
  29  ECA1228      0.26%    passed    100.00%
  30  ECA1229      0.26%    passed    100.00%
  31  ECA1232      0.26%    passed    100.00%
  32  ECA1237      0.26%    passed    100.00%
  33  ECA1238      0.26%    passed    100.00%
  34  ECA1240      0.26%    passed    100.00%
  35  ECA1242      0.26%    passed    100.00%
  36  ECA1243      0.26%    passed    100.00%
  37  ECA1247      0.26%    passed    100.00%
  38  ECA1251      0.26%    passed    100.00%
  39  ECA1253      0.26%    passed    100.00%
  40  ECA1255      0.26%    passed    100.00%
  41  ECA1257      0.26%    passed    100.00%
  42  ECA1260      0.26%    passed    100.00%
  43  ECA1261      0.26%    passed    100.00%
  44  ECA1271      0.26%    passed    100.00%
  45  ECA1276      0.26%    passed    100.00%
  46  ECA1278      0.26%    passed    100.00%
  47  ECA1281      0.26%    passed    100.00%
  48  ECA1283      0.26%    passed    100.00%
  49  ECA1284      0.26%    passed    100.00%
  50  ECA1286      0.26%    passed    100.00%
  51  ECA1287      0.26%    passed    100.00%
  52  ECA1288      0.26%    passed    100.00%
  53  ECA1289      0.26%    passed    100.00%
  54  ECA1293      0.26%    passed    100.00%
  55  ECA1298      0.26%    passed    100.00%
  56  ECA1316      0.26%    passed    100.00%
  57  ECA1409      0.26%    passed    100.00%
  58  ECA1413      0.26%    passed    100.00%
  59  ECA1493      0.26%    passed    100.00%
  60  ECA1693      0.26%    passed    100.00%
  61  ECA1713      0.26%    passed    100.00%
  62  ECA1717      0.26%    passed    100.00%
  63  ECA1725      0.26%    passed    100.00%
  64  ECA1751      0.26%    passed    100.00%
  65  ECA1757      0.26%    passed    100.00%
  66  ECA1761      0.26%    passed    100.00%
  67  ECA1769      0.26%    passed    100.00%
  68  ECA1793      0.26%    passed    100.00%
  69  ECA1805      0.26%    passed    100.00%
  70  ECA1815      0.26%    passed    100.00%
  71  ECA1821      0.26%    passed    100.00%
  72  ECA1825      0.26%    passed    100.00%
  73  ECA1851      0.26%    passed    100.00%
  74  ECA1853      0.26%    passed    100.00%
  75  ECA1875      0.26%    passed    100.00%
  76  ECA1885      0.26%    passed    100.00%
  77  ECA1889      0.26%    passed    100.00%
  78  ECA1891      0.26%    passed    100.00%
  79  ECA189       0.26%    passed    100.00%
  80  ECA1901      0.26%    passed    100.00%
  81  ECA1919      0.26%    passed    100.00%
  82  ECA191       0.26%    passed    100.00%
  83  ECA1939      0.26%    passed    100.00%
  84  ECA1969      0.26%    passed    100.00%
  85  ECA1997      0.26%    passed    100.00%
  86  ECA2041      0.26%    passed    100.00%
  87  ECA2043      0.26%    passed    100.00%
  88  ECA2073      0.26%    passed    100.00%
  89  ECA2081      0.26%    passed    100.00%
  90  ECA2091      0.26%    passed    100.00%
  91  ECA2111      0.26%    passed    100.00%
  92  ECA2151      0.26%    passed    100.00%
  93  ECA2191      0.26%    passed    100.00%
  94  ECA2195      0.26%    passed    100.00%
  95  ECA2199      0.26%    passed    100.00%
  96  ECA2251      0.26%    passed    100.00%
  97  ECA2291      0.26%    passed    100.00%
  98  ECA2332      0.26%    passed    100.00%
  99  ECA2334      0.26%    passed    100.00%
 100  ECA2336      0.26%    passed    100.00%
 101  ECA2350      0.26%    passed    100.00%
 102  ECA2360      0.26%    passed    100.00%
 103  ECA2367      0.26%    passed    100.00%
 104  ECA2377      0.26%    passed    100.00%
 105  ECA2417      0.26%    passed    100.00%
 106  ECA2437      0.26%    passed    100.00%
 107  ECA243       0.26%    passed    100.00%
 108  ECA2443      0.26%    passed    100.00%
 109  ECA2467      0.26%    passed    100.00%
 110  ECA246       0.26%    passed    100.00%
 111  ECA2473      0.26%    passed    100.00%
 112  ECA2475      0.26%    passed    100.00%
 113  ECA2481      0.26%    passed    100.00%
 114  ECA248       0.26%    passed    100.00%
 115  ECA2522      0.26%    passed    100.00%
 116  ECA2523      0.26%    passed    100.00%
 117  ECA2546      0.26%    passed    100.00%
 118  ECA2551      0.26%    passed    100.00%
 119  ECA2559      0.26%    passed    100.00%
 120  ECA2560      0.26%    passed    100.00%
 121  ECA2561      0.26%    passed    100.00%
 122  ECA2565      0.26%    passed    100.00%
 123  ECA2568      0.26%    passed    100.00%
 124  ECA2581      0.26%    passed    100.00%
 125  ECA2583      0.26%    passed    100.00%
 126  ECA2584      0.26%    passed    100.00%
 127  ECA259       0.26%    passed    100.00%
 128  ECA2601      0.26%    passed    100.00%
 129  ECA2602      0.26%    passed    100.00%
 130  ECA2607      0.26%    passed    100.00%
 131  ECA2609      0.26%    passed    100.00%
 132  ECA2648      0.26%    passed    100.00%
 133  ECA2656      0.26%    passed    100.00%
 134  ECA2660      0.26%    passed    100.00%
 135  ECA2676      0.26%    passed    100.00%
 136  ECA347       0.26%    passed    100.00%
 137  ECA363       0.26%    passed    100.00%
 138  ECA369       0.26%    passed    100.00%
 139  ECA372       0.26%    passed    100.00%
 140  ECA396       0.26%    passed    100.00%
 141  ECA593       0.26%    passed    100.00%
 142  ECA594       0.26%    passed    100.00%
 143  ECA701       0.26%    passed    100.00%
 144  ECA703       0.26%    passed    100.00%
 145  ECA705       0.26%    passed    100.00%
 146  ECA706       0.26%    passed    100.00%
 147  ECA710       0.26%    passed    100.00%
 148  ECA712       0.26%    passed    100.00%
 149  ECA722       0.26%    passed    100.00%
 150  ECA723       0.26%    passed    100.00%
 151  ECA724       0.26%    passed    100.00%
 152  ECA730       0.26%    passed    100.00%
 153  ECA732       0.26%    passed    100.00%
 154  ECA733       0.26%    passed    100.00%
 155  ECA738       0.26%    passed    100.00%
 156  ECA740       0.26%    passed    100.00%
 157  ECA741       0.26%    passed    100.00%
 158  ECA742       0.26%    passed    100.00%
 159  ECA743       0.26%    passed    100.00%
 160  ECA744       0.26%    passed    100.00%
 161  ECA745       0.26%    passed    100.00%
 162  ECA746       0.26%    passed    100.00%
 163  ECA760       0.26%    passed    100.00%
 164  ECA768       0.26%    passed    100.00%
 165  ECA923       0.26%    passed    100.00%
 166  ECA928       0.26%    passed    100.00%
 167  ED3011       0.26%    passed    100.00%
 168  ED3012       0.26%    passed    100.00%
 169  ED3040       0.26%    passed    100.00%
 170  ED3046       0.26%    passed    100.00%
 171  ED3052       0.26%    passed    100.00%
 172  ED3073       0.26%    passed    100.00%
 173  ED3077       0.26%    passed    100.00%
 174  EG4349       0.26%    passed    100.00%
 175  EG4724       0.26%    passed    100.00%
 176  EG4725       0.26%    passed    100.00%
 177  EG4946       0.26%    passed    100.00%
 178  JR4305       0.26%    passed    100.00%
 179  JU1088       0.26%    passed    100.00%
 180  JU1200       0.26%    passed    100.00%
 181  JU1246       0.26%    passed    100.00%
 182  JU1249       0.26%    passed    100.00%
 183  JU1395       0.26%    passed    100.00%
 184  JU1400       0.26%    passed    100.00%
 185  JU1409       0.26%    passed    100.00%
 186  JU1530       0.26%    passed    100.00%
 187  JU1543       0.26%    passed    100.00%
 188  JU1581       0.26%    passed    100.00%
 189  JU1792       0.26%    passed    100.00%
 190  JU1896       0.26%    passed    100.00%
 191  JU2001       0.26%    passed    100.00%
 192  JU2007       0.26%    passed    100.00%
 193  JU2016       0.26%    passed    100.00%
 194  JU2017       0.26%    passed    100.00%
 195  JU2131       0.26%    passed    100.00%
 196  JU2234       0.26%    passed    100.00%
 197  JU2250       0.26%    passed    100.00%
 198  JU2316       0.26%    passed    100.00%
 199  JU2464       0.26%    passed    100.00%
 200  JU2466       0.26%    passed    100.00%
 201  JU2513       0.26%    passed    100.00%
 202  JU2519       0.26%    passed    100.00%
 203  JU2526       0.26%    passed    100.00%
 204  JU2534       0.26%    passed    100.00%
 205  JU2575       0.26%    passed    100.00%
 206  JU2576       0.26%    passed    100.00%
 207  JU2578       0.26%    passed    100.00%
 208  JU2587       0.26%    passed    100.00%
 209  JU258        0.26%    passed    100.00%
 210  JU2592       0.26%    passed    100.00%
 211  JU2593       0.26%    passed    100.00%
 212  JU2800       0.26%    passed    100.00%
 213  JU2811       0.26%    passed    100.00%
 214  JU2825       0.26%    passed    100.00%
 215  JU2829       0.26%    passed    100.00%
 216  JU2841       0.26%    passed    100.00%
 217  JU2862       0.26%    passed    100.00%
 218  JU2866       0.26%    passed    100.00%
 219  JU2878       0.26%    passed    100.00%
 220  JU2879       0.26%    passed    100.00%
 221  JU2906       0.26%    passed    100.00%
 222  JU2907       0.26%    passed    100.00%
 223  JU310        0.26%    passed    100.00%
 224  JU3125       0.26%    passed    100.00%
 225  JU3127       0.26%    passed    100.00%
 226  JU3132       0.26%    passed    100.00%
 227  JU3134       0.26%    passed    100.00%
 228  JU3140       0.26%    passed    100.00%
 229  JU3144       0.26%    passed    100.00%
 230  JU3166       0.26%    passed    100.00%
 231  JU3167       0.26%    passed    100.00%
 232  JU3169       0.26%    passed    100.00%
 233  JU3224       0.26%    passed    100.00%
 234  JU3225       0.26%    passed    100.00%
 235  JU3226       0.26%    passed    100.00%
 236  JU323        0.26%    passed    100.00%
 237  JU3291       0.26%    passed    100.00%
 238  JU3318       0.26%    passed    100.00%
 239  JU360        0.26%    passed    100.00%
 240  JU3795       0.26%    passed    100.00%
 241  JU394        0.26%    passed    100.00%
 242  JU4047       0.26%    passed    100.00%
 243  JU406        0.26%    passed    100.00%
 244  JU4073       0.26%    passed    100.00%
 245  JU4161       0.26%    passed    100.00%
 246  JU561        0.26%    passed    100.00%
 247  JU642        0.26%    passed    100.00%
 248  JU782        0.26%    passed    100.00%
 249  JU830        0.26%    passed    100.00%
 250  LKC34        0.26%    passed    100.00%
 251  MY10         0.26%    passed    100.00%
 252  MY16         0.26%    passed    100.00%
 253  MY18         0.26%    passed    100.00%
 254  MY1          0.26%    passed    100.00%
 255  MY2147       0.26%    passed    100.00%
 256  MY23         0.26%    passed    100.00%
 257  MY2530       0.26%    passed    100.00%
 258  MY2573       0.26%    passed    100.00%
 259  MY2585       0.26%    passed    100.00%
 260  MY2693       0.26%    passed    100.00%
 261  MY2713       0.26%    passed    100.00%
 262  MY518        0.26%    passed    100.00%
 263  MY772        0.26%    passed    100.00%
 264  MY920        0.26%    passed    100.00%
 265  N2           0.26%    passed    100.00%
 266  NIC1049      0.26%    passed    100.00%
 267  NIC166       0.26%    passed    100.00%
 268  NIC1700      0.26%    passed    100.00%
 269  NIC1772      0.26%    passed    100.00%
 270  NIC1773      0.26%    passed    100.00%
 271  NIC1775      0.26%    passed    100.00%
 272  NIC1780      0.26%    passed    100.00%
 273  NIC1782      0.26%    passed    100.00%
 274  NIC1783      0.26%    passed    100.00%
 275  NIC1785      0.26%    passed    100.00%
 276  NIC1787      0.26%    passed    100.00%
 277  NIC1788      0.26%    passed    100.00%
 278  NIC1789      0.26%    passed    100.00%
 279  NIC1790      0.26%    passed    100.00%
 280  NIC1792      0.26%    passed    100.00%
 281  NIC1794      0.26%    passed    100.00%
 282  NIC1798      0.26%    passed    100.00%
 283  NIC1799      0.26%    passed    100.00%
 284  NIC1805      0.26%    passed    100.00%
 285  NIC1809      0.26%    passed    100.00%
 286  NIC1811      0.26%    passed    100.00%
 287  NIC199       0.26%    passed    100.00%
 288  NIC1         0.26%    passed    100.00%
 289  NIC207       0.26%    passed    100.00%
 290  NIC236       0.26%    passed    100.00%
 291  NIC251       0.26%    passed    100.00%
 292  NIC252       0.26%    passed    100.00%
 293  NIC259       0.26%    passed    100.00%
 294  NIC262       0.26%    passed    100.00%
 295  NIC266       0.26%    passed    100.00%
 296  NIC267       0.26%    passed    100.00%
 297  NIC268       0.26%    passed    100.00%
 298  NIC269       0.26%    passed    100.00%
 299  NIC274       0.26%    passed    100.00%
 300  NIC275       0.26%    passed    100.00%
 301  NIC277       0.26%    passed    100.00%
 302  NIC501       0.26%    passed    100.00%
 303  NIC511       0.26%    passed    100.00%
 304  NIC513       0.26%    passed    100.00%
 305  NIC523       0.26%    passed    100.00%
 306  NIC526       0.26%    passed    100.00%
 307  NIC527       0.26%    passed    100.00%
 308  NIC529       0.26%    passed    100.00%
 309  PB303        0.26%    passed    100.00%
 310  PX179        0.26%    passed    100.00%
 311  QG2075       0.26%    passed    100.00%
 312  QG2811       0.26%    passed    100.00%
 313  QG2813       0.26%    passed    100.00%
 314  QG2818       0.26%    passed    100.00%
 315  QG2825       0.26%    passed    100.00%
 316  QG2827       0.26%    passed    100.00%
 317  QG2828       0.26%    passed    100.00%
 318  QG2836       0.26%    passed    100.00%
 319  QG2837       0.26%    passed    100.00%
 320  QG2838       0.26%    passed    100.00%
 321  QG2843       0.26%    passed    100.00%
 322  QG2850       0.26%    passed    100.00%
 323  QG2855       0.26%    passed    100.00%
 324  QG2873       0.26%    passed    100.00%
 325  QG2874       0.26%    passed    100.00%
 326  QG2875       0.26%    passed    100.00%
 327  QG2877       0.26%    passed    100.00%
 328  QG2932       0.26%    passed    100.00%
 329  QG4003       0.26%    passed    100.00%
 330  QG4004       0.26%    passed    100.00%
 331  QG4010       0.26%    passed    100.00%
 332  QG4015       0.26%    passed    100.00%
 333  QG4017       0.26%    passed    100.00%
 334  QG4018       0.26%    passed    100.00%
 335  QG4080       0.26%    passed    100.00%
 336  QG4103       0.26%    passed    100.00%
 337  QG4135       0.26%    passed    100.00%
 338  QG4138       0.26%    passed    100.00%
 339  QG4147       0.26%    passed    100.00%
 340  QG4151       0.26%    passed    100.00%
 341  QG4226       0.26%    passed    100.00%
 342  QG4228       0.26%    passed    100.00%
 343  QG536        0.26%    passed    100.00%
 344  QG556        0.26%    passed    100.00%
 345  QG557        0.26%    passed    100.00%
 346  QX1211       0.26%    passed    100.00%
 347  QX1212       0.26%    passed    100.00%
 348  QX1791       0.26%    passed    100.00%
 349  QX1792       0.26%    passed    100.00%
 350  QX1793       0.26%    passed    100.00%
 351  QX1794       0.26%    passed    100.00%
 352  WN2001       0.26%    passed    100.00%
 353  WN2002       0.26%    passed    100.00%
 354  WN2063       0.26%    passed    100.00%
 355  WN2066       0.26%    passed    100.00%
 356  WN2068       0.26%    passed    100.00%
 357  WN2069       0.26%    passed    100.00%
 358  WN2086       0.26%    passed    100.00%
 359  WN2117       0.26%    passed    100.00%
 360  XZ1515       0.26%    passed    100.00%
 361  XZ1516       0.26%    passed    100.00%
 362  XZ1672       0.26%    passed    100.00%
 363  XZ1734       0.26%    passed    100.00%
 364  XZ1735       0.26%    passed    100.00%
 365  XZ1756       0.26%    passed    100.00%
 366  XZ2019       0.26%    passed    100.00%
 367  XZ2020       0.26%    passed    100.00%
 368  XZ2211       0.26%    passed    100.00%
****  TOTAL        0.26%  0 sequences failed composition chi2 test (p-value<5%; df=19)
Reading constraint tree constraint...
INFO: Constraint tree will be applied to ML tree and all bootstrap trees.
NOTE: BRC20067 is identical to AB1 but kept for subsequent analysis
NOTE: CX11314 is identical to CX11254 but kept for subsequent analysis
NOTE: ECA2602 is identical to CX11307 but kept for subsequent analysis
NOTE: ECA928 is identical to DL226 but kept for subsequent analysis
NOTE: ED3040 is identical to ECA1069 but kept for subsequent analysis
NOTE: ECA1413 is identical to ECA1200 but kept for subsequent analysis
NOTE: ECA2073 is identical to ECA1202 but kept for subsequent analysis
NOTE: ECA1997 is identical to ECA1206 but kept for subsequent analysis
NOTE: ECA2251 is identical to ECA1247 but kept for subsequent analysis
NOTE: ECA2481 is identical to ECA1271 but kept for subsequent analysis
NOTE: ECA1891 is identical to ECA1286 but kept for subsequent analysis
NOTE: ECA2377 is identical to ECA1288 but kept for subsequent analysis
NOTE: ECA1751 is identical to ECA1409 but kept for subsequent analysis
NOTE: ECA1761 is identical to ECA1725 but kept for subsequent analysis
NOTE: ECA1853 is identical to ECA1815 but kept for subsequent analysis
NOTE: ECA2336 is identical to ECA1825 but kept for subsequent analysis
NOTE: ECA1885 is identical to ECA1851 but kept for subsequent analysis
NOTE: ECA2332 is identical to ECA1875 but kept for subsequent analysis
NOTE: ECA744 is identical to ECA1969 but kept for subsequent analysis
NOTE: MY16 is identical to ECA2523 but kept for subsequent analysis
NOTE: JU2841 is identical to ECA2546 but kept for subsequent analysis
NOTE: JU1200 is identical to ECA2551 but kept for subsequent analysis
NOTE: JU3226 is identical to ECA2581 but kept for subsequent analysis
NOTE: EG4724 is identical to ECA594 but kept for subsequent analysis
NOTE: ECA712 is identical to ECA710 but kept for subsequent analysis
NOTE: JU2234 is identical to JU1246 but kept for subsequent analysis
NOTE: JU2007 is identical to JU1581 but kept for subsequent analysis
NOTE: XZ2020 is identical to JU2001 but kept for subsequent analysis
NOTE: JU2316 is identical to JU2131 but kept for subsequent analysis
NOTE: JU830 is identical to JU2575 but kept for subsequent analysis
NOTE: NIC1794 is identical to NIC1788 but kept for subsequent analysis
NOTE: NIC1798 is identical to NIC1790 but kept for subsequent analysis
NOTE: WN2086 is identical to PB303 but kept for subsequent analysis
NOTE: QG2874 is identical to QG2873 but kept for subsequent analysis
NOTE: 95 identical sequences (see below) will be ignored for subsequent analysis
NOTE: CB4852 (identical to AB1) is ignored but added at the end
NOTE: CX11271 (identical to AB1) is ignored but added at the end
NOTE: ECA2559 (identical to AB1) is ignored but added at the end
NOTE: ECA2568 (identical to AB1) is ignored but added at the end
NOTE: ECA2660 (identical to AB1) is ignored but added at the end
NOTE: ECA923 (identical to AB1) is ignored but added at the end
NOTE: ED3052 (identical to AB1) is ignored but added at the end
NOTE: ED3077 (identical to AB1) is ignored but added at the end
NOTE: JU1395 (identical to AB1) is ignored but added at the end
NOTE: JU1400 (identical to AB1) is ignored but added at the end
NOTE: JU1530 (identical to AB1) is ignored but added at the end
NOTE: JU2016 (identical to AB1) is ignored but added at the end
NOTE: JU2250 (identical to AB1) is ignored but added at the end
NOTE: JU2464 (identical to AB1) is ignored but added at the end
NOTE: JU2534 (identical to AB1) is ignored but added at the end
NOTE: JU2829 (identical to AB1) is ignored but added at the end
NOTE: JU2862 (identical to AB1) is ignored but added at the end
NOTE: JU2878 (identical to AB1) is ignored but added at the end
NOTE: JU3134 (identical to AB1) is ignored but added at the end
NOTE: JU3140 (identical to AB1) is ignored but added at the end
NOTE: JU3144 (identical to AB1) is ignored but added at the end
NOTE: JU3224 (identical to AB1) is ignored but added at the end
NOTE: JU4073 (identical to AB1) is ignored but added at the end
NOTE: JU642 (identical to AB1) is ignored but added at the end
NOTE: MY1 (identical to AB1) is ignored but added at the end
NOTE: MY2573 (identical to AB1) is ignored but added at the end
NOTE: MY518 (identical to AB1) is ignored but added at the end
NOTE: NIC1049 (identical to AB1) is ignored but added at the end
NOTE: NIC1772 (identical to AB1) is ignored but added at the end
NOTE: NIC1805 (identical to AB1) is ignored but added at the end
NOTE: NIC1811 (identical to AB1) is ignored but added at the end
NOTE: NIC207 (identical to AB1) is ignored but added at the end
NOTE: NIC526 (identical to AB1) is ignored but added at the end
NOTE: NIC529 (identical to AB1) is ignored but added at the end
NOTE: QG2818 (identical to AB1) is ignored but added at the end
NOTE: QG2836 (identical to AB1) is ignored but added at the end
NOTE: QG2843 (identical to AB1) is ignored but added at the end
NOTE: QG2850 (identical to AB1) is ignored but added at the end
NOTE: QG2855 (identical to AB1) is ignored but added at the end
NOTE: QG4010 (identical to AB1) is ignored but added at the end
NOTE: QG4017 (identical to AB1) is ignored but added at the end
NOTE: QG4138 (identical to AB1) is ignored but added at the end
NOTE: QG4151 (identical to AB1) is ignored but added at the end
NOTE: QG536 (identical to AB1) is ignored but added at the end
NOTE: QX1212 (identical to AB1) is ignored but added at the end
NOTE: WN2001 (identical to AB1) is ignored but added at the end
NOTE: WN2069 (identical to AB1) is ignored but added at the end
NOTE: XZ1672 (identical to AB1) is ignored but added at the end
NOTE: XZ1756 (identical to AB1) is ignored but added at the end
NOTE: ECA705 (identical to CX11307) is ignored but added at the end
NOTE: QG2813 (identical to CX11307) is ignored but added at the end
NOTE: QG2838 (identical to CX11307) is ignored but added at the end
NOTE: MY2530 (identical to ECA1069) is ignored but added at the end
NOTE: NIC1782 (identical to ECA1069) is ignored but added at the end
NOTE: ECA2473 (identical to ECA1206) is ignored but added at the end
NOTE: ECA732 (identical to ECA1288) is ignored but added at the end
NOTE: ECA2151 (identical to ECA1409) is ignored but added at the end
NOTE: ECA2350 (identical to ECA1815) is ignored but added at the end
NOTE: QG557 (identical to ECA1815) is ignored but added at the end
NOTE: ECA745 (identical to ECA1969) is ignored but added at the end
NOTE: JU4161 (identical to ECA2546) is ignored but added at the end
NOTE: JU2592 (identical to ECA2551) is ignored but added at the end
NOTE: JU2593 (identical to ECA2551) is ignored but added at the end
NOTE: JU394 (identical to ECA2551) is ignored but added at the end
NOTE: JU561 (identical to ECA2551) is ignored but added at the end
NOTE: N2 (identical to ECA2551) is ignored but added at the end
NOTE: QG2811 (identical to ECA2551) is ignored but added at the end
NOTE: QG2825 (identical to ECA2551) is ignored but added at the end
NOTE: QG2837 (identical to ECA2551) is ignored but added at the end
NOTE: QG4004 (identical to ECA2551) is ignored but added at the end
NOTE: QG4080 (identical to ECA2551) is ignored but added at the end
NOTE: QG4228 (identical to ECA2551) is ignored but added at the end
NOTE: NIC199 (identical to ECA594) is ignored but added at the end
NOTE: NIC268 (identical to ECA594) is ignored but added at the end
NOTE: NIC274 (identical to ECA594) is ignored but added at the end
NOTE: NIC513 (identical to ECA594) is ignored but added at the end
NOTE: ECA768 (identical to ECA710) is ignored but added at the end
NOTE: JU2906 (identical to JU1246) is ignored but added at the end
NOTE: JU782 (identical to JU1246) is ignored but added at the end
NOTE: MY10 (identical to JU1246) is ignored but added at the end
NOTE: MY2147 (identical to JU1246) is ignored but added at the end
NOTE: MY2693 (identical to JU1246) is ignored but added at the end
NOTE: MY2713 (identical to JU1246) is ignored but added at the end
NOTE: NIC1773 (identical to JU1246) is ignored but added at the end
NOTE: NIC1 (identical to JU1246) is ignored but added at the end
NOTE: NIC251 (identical to JU1246) is ignored but added at the end
NOTE: NIC511 (identical to JU1246) is ignored but added at the end
NOTE: WN2117 (identical to JU1246) is ignored but added at the end
NOTE: NIC166 (identical to JU1581) is ignored but added at the end
NOTE: NIC527 (identical to JU1581) is ignored but added at the end
NOTE: JU2526 (identical to JU2131) is ignored but added at the end
NOTE: JU3127 (identical to JU2131) is ignored but added at the end
NOTE: NIC1785 (identical to JU2131) is ignored but added at the end
NOTE: NIC1789 (identical to JU2131) is ignored but added at the end
NOTE: NIC262 (identical to JU2131) is ignored but added at the end
95 taxa removed from constraint tree

For your convenience alignment with unique sequences printed to FINAL.fasta.uniqueseq.phy

Creating fast initial parsimony tree by random order stepwise addition...
0.012 seconds, parsimony score: 807 (based on 600 sites)
NOTE: ModelFinder requires 185 MB RAM!
ModelFinder will test 546 protein models (sample size: 3436) ...
 No. Model         -LnL         df  AIC          AICc         BIC
Measuring multi-threading efficiency up to 48 CPU cores
Increase to 10 rounds for branch lengths
157 trees examined
Threads: 1 / Time: 48.090 sec / Speedup: 1.000 / Efficiency: 100% / LogL: -25434
Threads: 2 / Time: 28.321 sec / Speedup: 1.698 / Efficiency: 85% / LogL: -25434
Threads: 3 / Time: 21.957 sec / Speedup: 2.190 / Efficiency: 73% / LogL: -25434
Threads: 4 / Time: 18.838 sec / Speedup: 2.553 / Efficiency: 64% / LogL: -25434
Threads: 5 / Time: 17.337 sec / Speedup: 2.774 / Efficiency: 55% / LogL: -25434
Threads: 6 / Time: 15.838 sec / Speedup: 3.036 / Efficiency: 51% / LogL: -25434
Threads: 7 / Time: 15.226 sec / Speedup: 3.158 / Efficiency: 45% / LogL: -25434
BEST NUMBER OF THREADS: 6

  1  Dayhoff       16752.682    543 34591.363    34795.646    37926.504
  2  Dayhoff+I     16610.580    544 34309.160    34514.265    37650.442
  3  Dayhoff+G4    16599.015    544 34286.031    34491.136    37627.313
  4  Dayhoff+I+G4  16588.967    545 34267.934    34473.865    37615.359
  5  Dayhoff+R2    16567.482    545 34224.963    34430.894    37572.388
  6  Dayhoff+R3    16563.874    547 34221.748    34429.335    37581.456
 14  Dayhoff+F     15553.291    562 32230.582    32450.844    35682.422
 15  Dayhoff+F+I   15428.867    563 31983.734    32204.856    35441.715
 16  Dayhoff+F+G4  15417.926    563 31961.853    32182.975    35419.834
 17  Dayhoff+F+I+G4 15409.268    564 31946.535    32168.521    35410.659
 18  Dayhoff+F+R2  15387.722    564 31903.443    32125.429    35367.567
 19  Dayhoff+F+R3  15372.467    566 31876.934    32100.651    35353.342
 20  Dayhoff+F+R4  15366.664    568 31869.329    32094.785    35358.021
 27  mtMAM         15986.913    543 33059.826    33264.108    36394.967
 28  mtMAM+I       15833.816    544 32755.631    32960.737    36096.914
 29  mtMAM+G4      15817.128    544 32722.257    32927.362    36063.539
 30  mtMAM+I+G4    15806.476    545 32702.953    32908.883    36050.377
 31  mtMAM+R2      15794.739    545 32679.477    32885.408    36026.902
 32  mtMAM+R3      15773.147    547 32640.294    32847.881    36000.003
 33  mtMAM+R4      15763.506    549 32625.011    32834.263    35997.004
 34  mtMAM+R5      15750.767    551 32603.535    32814.459    35987.812
 35  mtMAM+R6      15745.968    553 32597.936    32810.540    35994.497
 40  mtMAM+F       15561.925    562 32247.849    32468.111    35699.689
 41  mtMAM+F+I     15401.018    563 31928.036    32149.158    35386.017
 42  mtMAM+F+G4    15386.370    563 31898.740    32119.862    35356.722
 43  mtMAM+F+I+G4  15374.996    564 31877.992    32099.977    35342.115
 44  mtMAM+F+R2    15363.470    564 31854.940    32076.925    35319.064
 45  mtMAM+F+R3    15343.328    566 31818.655    32042.372    35295.063
 46  mtMAM+F+R4    15333.657    568 31803.314    32028.771    35292.006
 47  mtMAM+F+R5    15321.152    570 31782.303    32009.507    35283.279
 48  mtMAM+F+R6    15316.457    572 31776.914    32005.874    35290.174
 53  JTT           16380.156    543 33846.313    34050.595    37181.453
 54  JTT+I         16259.083    544 33606.167    33811.272    36947.449
 55  JTT+G4        16248.960    544 33585.920    33791.025    36927.202
 56  JTT+I+G4      16240.963    545 33571.926    33777.857    36919.351
 57  JTT+R2        16222.560    545 33535.120    33741.050    36882.544
 58  JTT+R3        16208.459    547 33510.917    33718.504    36870.626
 59  JTT+R4        16202.599    549 33503.199    33712.450    36875.192
 66  JTT+F         15439.585    562 32003.169    32223.431    35455.009
 67  JTT+F+I       15325.714    563 31777.428    31998.550    35235.409
 68  JTT+F+G4      15315.273    563 31756.546    31977.668    35214.527
 69  JTT+F+I+G4    15307.828    564 31743.656    31965.641    35207.779
 70  JTT+F+R2      15289.259    564 31706.517    31928.503    35170.641
 71  JTT+F+R3      15273.285    566 31678.569    31902.286    35154.977
 72  JTT+F+R4      15266.877    568 31669.755    31895.211    35158.447
 79  WAG           16532.484    543 34150.969    34355.251    37486.109
 80  WAG+I         16407.321    544 33902.642    34107.747    37243.924
 81  WAG+G4        16397.476    544 33882.951    34088.057    37224.234
 82  WAG+I+G4      16388.797    545 33867.594    34073.524    37215.018
 83  WAG+R2        16368.205    545 33826.409    34032.340    37173.834
 84  WAG+R3        16354.252    547 33802.505    34010.092    37162.213
 85  WAG+R4        16348.000    549 33794.000    34003.252    37165.993
 92  WAG+F         15531.271    562 32186.543    32406.804    35638.382
 93  WAG+F+I       15412.043    563 31950.087    32171.209    35408.068
 94  WAG+F+G4      15401.677    563 31929.354    32150.476    35387.335
 95  WAG+F+I+G4    15393.302    564 31914.604    32136.590    35378.728
 96  WAG+F+R2      15371.909    564 31871.819    32093.804    35335.943
 97  WAG+F+R3      15356.413    566 31844.827    32068.543    35321.234
 98  WAG+F+R4      15349.565    568 31835.130    32060.587    35323.822
105  cpREV         16332.083    543 33750.165    33954.447    37085.306
106  cpREV+I       16195.685    544 33479.369    33684.475    36820.652
107  cpREV+G4      16186.148    544 33460.296    33665.402    36801.579
108  cpREV+I+G4    16176.908    545 33443.815    33649.746    36791.240
109  cpREV+R2      16160.399    545 33410.799    33616.730    36758.223
110  cpREV+R3      16146.710    547 33387.419    33595.007    36747.128
111  cpREV+R4      16140.316    549 33378.631    33587.883    36750.624
118  cpREV+F       15541.555    562 32207.111    32427.372    35658.950
119  cpREV+F+I     15419.021    563 31964.042    32185.164    35422.024
120  cpREV+F+G4    15409.089    563 31944.179    32165.301    35402.160
121  cpREV+F+I+G4  15400.640    564 31929.280    32151.266    35393.404
122  cpREV+F+R2    15381.363    564 31890.726    32112.711    35354.849
123  cpREV+F+R3    15366.478    566 31864.957    32088.674    35341.364
124  cpREV+F+R4    15359.849    568 31855.698    32081.155    35344.390
131  mtREV         15973.910    543 33033.821    33238.103    36368.961
132  mtREV+I       15849.099    544 32786.198    32991.303    36127.480
133  mtREV+G4      15836.361    544 32760.722    32965.827    36102.004
134  mtREV+I+G4    15827.695    545 32745.390    32951.321    36092.814
135  mtREV+R2      15808.443    545 32706.886    32912.817    36054.310
136  mtREV+R3      15791.853    547 32677.705    32885.293    36037.414
137  mtREV+R4      15784.740    549 32667.480    32876.732    36039.473
144  mtREV+F       15507.284    562 32138.567    32358.829    35590.407
145  mtREV+F+I     15381.994    563 31889.988    32111.111    35347.970
146  mtREV+F+G4    15369.245    563 31864.489    32085.612    35322.471
147  mtREV+F+I+G4  15360.246    564 31848.492    32070.477    35312.615
148  mtREV+F+R2    15339.896    564 31807.791    32029.777    35271.915
149  mtREV+F+R3    15323.488    566 31778.975    32002.692    35255.383
150  mtREV+F+R4    15316.302    568 31768.604    31994.061    35257.296
157  rtREV         16867.130    543 34820.259    35024.542    38155.400
158  rtREV+I       16734.335    544 34556.670    34761.775    37897.952
159  rtREV+G4      16724.463    544 34536.927    34742.032    37878.209
160  rtREV+I+G4    16715.239    545 34520.477    34726.408    37867.902
161  rtREV+R2      16698.867    545 34487.733    34693.664    37835.158
162  rtREV+R3      16683.238    547 34460.477    34668.064    37820.186
163  rtREV+R4      16676.262    549 34450.525    34659.777    37822.518
170  rtREV+F       15658.013    562 32440.026    32660.288    35891.866
171  rtREV+F+I     15520.569    563 32167.138    32388.260    35625.119
172  rtREV+F+G4    15510.925    563 32147.851    32368.974    35605.833
173  rtREV+F+I+G4  15501.417    564 32130.835    32352.820    35594.959
174  rtREV+F+R2    15487.576    564 32103.153    32325.138    35567.276
175  rtREV+F+R3    15470.554    566 32073.108    32296.825    35549.515
176  rtREV+F+R4    15462.687    568 32061.375    32286.831    35550.067
183  mtART         15712.203    543 32510.406    32714.688    35845.546
184  mtART+I       15577.215    544 32242.430    32447.535    35583.712
185  mtART+G4      15563.732    544 32215.464    32420.570    35556.747
186  mtART+I+G4    15554.348    545 32198.697    32404.627    35546.121
187  mtART+R2      15539.183    545 32168.366    32374.297    35515.791
188  mtART+R3      15520.960    547 32135.921    32343.508    35495.629
189  mtART+R4      15512.505    549 32123.009    32332.261    35495.002
190  mtART+R5      15502.704    551 32107.408    32318.332    35491.685
191  mtART+R6      15499.072    553 32104.144    32316.748    35500.705
196  mtART+F       15583.996    562 32291.992    32512.253    35743.831
197  mtART+F+I     15447.476    563 32020.953    32242.076    35478.935
198  mtART+F+G4    15433.780    563 31993.559    32214.682    35451.541
199  mtART+F+I+G4  15423.983    564 31975.966    32197.952    35440.090
200  mtART+F+R2    15407.774    564 31943.547    32165.532    35407.671
201  mtART+F+R3    15389.872    566 31911.743    32135.460    35388.151
202  mtART+F+R4    15381.393    568 31898.785    32124.242    35387.477
203  mtART+F+R5    15371.619    570 31883.239    32110.443    35384.215
204  mtART+F+R6    15367.995    572 31879.990    32108.950    35393.250
209  mtZOA         15710.855    543 32507.710    32711.992    35842.850
210  mtZOA+I       15592.434    544 32272.869    32477.974    35614.151
211  mtZOA+G4      15578.848    544 32245.696    32450.801    35586.978
212  mtZOA+I+G4    15570.795    545 32231.590    32437.521    35579.015
213  mtZOA+R2      15552.574    545 32195.148    32401.078    35542.572
214  mtZOA+R3      15534.832    547 32163.664    32371.252    35523.373
215  mtZOA+R4      15527.316    549 32152.633    32361.884    35524.625
222  mtZOA+F       15492.651    562 32109.301    32329.563    35561.141
223  mtZOA+F+I     15373.307    563 31872.614    32093.737    35330.596
224  mtZOA+F+G4    15360.271    563 31846.543    32067.665    35304.525
225  mtZOA+F+I+G4  15352.095    564 31832.191    32054.176    35296.314
226  mtZOA+F+R2    15334.207    564 31796.414    32018.399    35260.538
227  mtZOA+F+R3    15316.475    566 31764.950    31988.667    35241.358
228  mtZOA+F+R4    15308.857    568 31753.715    31979.171    35242.406
235  VT            16427.691    543 33941.382    34145.664    37276.522
236  VT+I          16312.899    544 33713.798    33918.903    37055.080
237  VT+G4         16303.196    544 33694.391    33899.497    37035.674
238  VT+I+G4       16295.360    545 33680.719    33886.650    37028.144
239  VT+R2         16275.024    545 33640.047    33845.978    36987.472
240  VT+R3         16260.107    547 33614.214    33821.801    36973.922
241  VT+R4         16253.693    549 33605.387    33814.638    36977.379
248  VT+F          15528.039    562 32180.077    32400.339    35631.917
249  VT+F+I        15417.207    563 31960.414    32181.537    35418.396
250  VT+F+G4       15407.032    563 31940.064    32161.187    35398.046
251  VT+F+I+G4     15399.450    564 31926.900    32148.886    35391.024
252  VT+F+R2       15378.646    564 31885.292    32107.277    35349.416
253  VT+F+R3       15362.345    566 31856.690    32080.407    35333.098
254  VT+F+R4       15355.477    568 31846.953    32072.410    35335.645
261  LG            16448.010    543 33982.021    34186.303    37317.161
262  LG+I          16322.678    544 33733.356    33938.461    37074.638
263  LG+G4         16312.620    544 33713.240    33918.345    37054.522
264  LG+I+G4       16303.766    545 33697.532    33903.463    37044.957
265  LG+R2         16285.224    545 33660.448    33866.379    37007.873
266  LG+R3         16269.996    547 33633.992    33841.579    36993.700
267  LG+R4         16263.243    549 33624.485    33833.737    36996.478
274  LG+F          15552.403    562 32228.807    32449.069    35680.646
275  LG+F+I        15425.503    563 31977.006    32198.129    35434.988
276  LG+F+G4       15415.159    563 31956.318    32177.441    35414.300
277  LG+F+I+G4     15406.116    564 31940.231    32162.217    35404.355
278  LG+F+R2       15388.669    564 31905.338    32127.323    35369.462
279  LG+F+R3       15371.983    566 31875.966    32099.683    35352.374
280  LG+F+R4       15364.419    568 31864.838    32090.295    35353.530
287  DCMut         16751.661    543 34589.321    34793.603    37924.462
288  DCMut+I       16609.912    544 34307.824    34512.930    37649.107
289  DCMut+G4      16598.405    544 34284.810    34489.916    37626.093
290  DCMut+I+G4    16588.006    545 34266.012    34471.943    37613.437
291  DCMut+R2      16569.101    545 34228.202    34434.133    37575.626
292  DCMut+R3      16554.232    547 34202.464    34410.051    37562.173
293  DCMut+R4      16547.636    549 34193.272    34402.524    37565.265
300  DCMut+F       15553.602    562 32231.205    32451.467    35683.045
301  DCMut+F+I     15429.280    563 31984.561    32205.683    35442.543
302  DCMut+F+G4    15418.354    563 31962.709    32183.831    35420.690
303  DCMut+F+I+G4  15409.418    564 31946.836    32168.821    35410.959
304  DCMut+F+R2    15388.949    564 31905.898    32127.884    35370.022
305  DCMut+F+R3    15372.615    566 31877.229    32100.946    35353.637
306  DCMut+F+R4    15365.407    568 31866.815    32092.271    35355.507
313  PMB           16422.615    543 33931.231    34135.513    37266.371
314  PMB+I         16309.850    544 33707.700    33912.806    37048.983
315  PMB+G4        16299.928    544 33687.855    33892.961    37029.138
316  PMB+I+G4      16292.024    545 33674.048    33879.979    37021.473
317  PMB+R2        16269.546    545 33629.093    33835.024    36976.517
318  PMB+R3        16254.486    547 33602.973    33810.560    36962.682
319  PMB+R4        16247.763    549 33593.526    33802.778    36965.519
326  PMB+F         15615.714    562 32355.428    32575.689    35807.267
327  PMB+F+I       15501.211    563 32128.421    32349.544    35586.403
328  PMB+F+G4      15490.815    563 32107.629    32328.752    35565.611
329  PMB+F+I+G4    15482.710    564 32093.419    32315.404    35557.543
330  PMB+F+R2      15460.829    564 32049.659    32271.644    35513.783
331  PMB+F+R3      15444.970    566 32021.939    32245.656    35498.347
332  PMB+F+R4      15437.744    568 32011.487    32236.944    35500.179
339  HIVb          16666.004    543 34418.007    34622.289    37753.147
340  HIVb+I        16503.602    544 34095.205    34300.310    37436.487
341  HIVb+G4       16495.662    544 34079.323    34284.429    37420.606
342  HIVb+I+G4     16485.713    545 34061.427    34267.358    37408.851
343  HIVb+R2       16477.443    545 34044.887    34250.817    37392.311
344  HIVb+R3       16464.156    547 34022.312    34229.899    37382.021
345  HIVb+R4       16457.608    549 34013.216    34222.467    37385.208
352  HIVb+F        15523.873    562 32171.747    32392.009    35623.586
353  HIVb+F+I      15356.898    563 31839.796    32060.918    35297.778
354  HIVb+F+G4     15348.652    563 31823.304    32044.426    35281.285
355  HIVb+F+I+G4   15338.050    564 31804.100    32026.086    35268.224
356  HIVb+F+R2     15329.645    564 31787.290    32009.276    35251.414
357  HIVb+F+R3     15316.214    566 31764.428    31988.145    35240.836
358  HIVb+F+R4     15308.838    568 31753.677    31979.133    35242.369
365  HIVw          16843.818    543 34773.637    34977.919    38108.777
366  HIVw+I        16655.029    544 34398.058    34603.164    37739.341
367  HIVw+G4       16647.524    544 34383.048    34588.153    37724.330
368  HIVw+I+G4     16636.752    545 34363.505    34569.435    37710.929
369  HIVw+R2       16631.783    545 34353.567    34559.497    37700.991
370  HIVw+R3       16619.563    547 34333.126    34540.713    37692.835
371  HIVw+R4       16612.300    549 34322.601    34531.852    37694.593
378  HIVw+F        15541.294    562 32206.588    32426.850    35658.428
379  HIVw+F+I      15379.352    563 31884.703    32105.826    35342.685
380  HIVw+F+G4     15369.661    563 31865.321    32086.444    35323.303
381  HIVw+F+I+G4   15359.229    564 31846.458    32068.443    35310.582
382  HIVw+F+R2     15346.770    564 31821.540    32043.525    35285.664
383  HIVw+F+R3     15333.582    566 31799.163    32022.880    35275.571
384  HIVw+F+R4     15326.106    568 31788.211    32013.668    35276.903
391  JTTDCMut      16394.794    543 33875.589    34079.871    37210.729
392  JTTDCMut+I    16273.173    544 33634.345    33839.451    36975.628
393  JTTDCMut+G4   16263.027    544 33614.054    33819.159    36955.336
394  JTTDCMut+I+G4 16254.698    545 33599.395    33805.326    36946.819
395  JTTDCMut+R2   16236.209    545 33562.418    33768.349    36909.842
396  JTTDCMut+R3   16222.199    547 33538.398    33745.985    36898.106
397  JTTDCMut+R4   16215.628    549 33529.256    33738.508    36901.249
404  JTTDCMut+F    15447.802    562 32019.605    32239.867    35471.444
405  JTTDCMut+F+I  15333.374    563 31792.748    32013.871    35250.730
406  JTTDCMut+F+G4 15322.966    563 31771.933    31993.055    35229.914
407  JTTDCMut+F+I+G4 15315.267    564 31758.534    31980.520    35222.658
408  JTTDCMut+F+R2 15296.780    564 31721.559    31943.545    35185.683
409  JTTDCMut+F+R3 15281.160    566 31694.320    31918.037    35170.728
410  JTTDCMut+F+R4 15274.045    568 31684.089    31909.546    35172.781
417  FLU           16510.030    543 34106.060    34310.342    37441.201
418  FLU+I         16362.660    544 33813.321    34018.426    37154.603
419  FLU+G4        16351.669    544 33791.338    33996.444    37132.621
420  FLU+I+G4      16341.756    545 33773.512    33979.443    37120.936
421  FLU+R2        16328.796    545 33747.591    33953.522    37095.016
422  FLU+R3        16314.566    547 33723.131    33930.718    37082.840
423  FLU+R4        16306.973    549 33711.947    33921.198    37083.940
430  FLU+F         15512.658    562 32149.316    32369.577    35601.155
431  FLU+F+I       15363.731    563 31853.461    32074.584    35311.443
432  FLU+F+G4      15349.770    563 31825.540    32046.663    35283.522
433  FLU+F+I+G4    15338.933    564 31805.866    32027.851    35269.989
434  FLU+F+R2      15324.483    564 31776.967    31998.952    35241.090
435  FLU+F+R3      15309.277    566 31750.555    31974.272    35226.962
436  FLU+F+R4      15300.943    568 31737.885    31963.342    35226.577
437  FLU+F+R5      15291.544    570 31723.088    31950.292    35224.064
438  FLU+F+R6      15288.115    572 31720.230    31949.190    35233.490
443  Blosum62      16418.782    543 33923.564    34127.846    37258.704
444  Blosum62+I    16303.903    544 33695.807    33900.912    37037.089
445  Blosum62+G4   16293.982    544 33675.964    33881.070    37017.247
446  Blosum62+I+G4 16285.852    545 33661.705    33867.636    37009.129
447  Blosum62+R2   16264.178    545 33618.355    33824.286    36965.780
448  Blosum62+R3   16249.491    547 33592.983    33800.570    36952.691
449  Blosum62+R4   16242.348    549 33582.695    33791.947    36954.688
456  Blosum62+F    15607.069    562 32338.138    32558.400    35789.978
457  Blosum62+F+I  15492.731    563 32111.461    32332.584    35569.443
458  Blosum62+F+G4 15482.461    563 32090.922    32312.044    35548.903
459  Blosum62+F+I+G4 15474.361    564 32076.722    32298.708    35540.846
460  Blosum62+F+R2 15453.272    564 32034.543    32256.529    35498.667
461  Blosum62+F+R3 15437.816    566 32007.632    32231.349    35484.040
462  Blosum62+F+R4 15430.238    568 31996.476    32221.933    35485.168
469  mtMet         15522.982    543 32131.965    32336.247    35467.105
470  mtMet+I       15412.326    544 31912.651    32117.757    35253.934
471  mtMet+G4      15399.907    544 31887.814    32092.920    35229.097
472  mtMet+I+G4    15392.849    545 31875.697    32081.628    35223.122
473  mtMet+R2      15376.032    545 31842.065    32047.995    35189.489
474  mtMet+R3      15358.580    547 31811.160    32018.748    35170.869
475  mtMet+R4      15350.903    549 31799.806    32009.058    35171.799
482  mtMet+F       15394.961    562 31913.922    32134.184    35365.762
483  mtMet+F+I     15282.654    563 31691.307    31912.430    35149.289
484  mtMet+F+G4    15270.381    563 31666.762    31887.885    35124.744
485  mtMet+F+I+G4  15263.072    564 31654.145    31876.130    35118.268
486  mtMet+F+R2    15245.808    564 31619.617    31841.602    35083.741
487  mtMet+F+R3    15228.684    566 31589.368    31813.085    35065.776
488  mtMet+F+R4    15220.946    568 31577.892    31803.349    35066.584
495  mtVer         15981.440    543 33048.881    33253.163    36384.021
496  mtVer+I       15854.643    544 32797.286    33002.391    36138.568
497  mtVer+G4      15841.186    544 32770.372    32975.477    36111.654
498  mtVer+I+G4    15833.035    545 32756.070    32962.001    36103.495
499  mtVer+R2      15821.091    545 32732.182    32938.113    36079.607
500  mtVer+R3      15804.022    547 32702.043    32909.631    36061.752
501  mtVer+R4      15795.947    549 32689.895    32899.146    36061.888
508  mtVer+F       15448.012    562 32020.024    32240.286    35471.863
509  mtVer+F+I     15313.338    563 31752.677    31973.799    35210.658
510  mtVer+F+G4    15302.079    563 31730.159    31951.281    35188.140
511  mtVer+F+I+G4  15293.149    564 31714.298    31936.283    35178.421
512  mtVer+F+R2    15282.136    564 31692.272    31914.258    35156.396
513  mtVer+F+R3    15266.225    566 31664.450    31888.167    35140.857
514  mtVer+F+R4    15258.101    568 31652.202    31877.659    35140.894
521  mtInv         15494.190    543 32074.380    32278.662    35409.520
522  mtInv+I       15385.426    544 31858.851    32063.957    35200.134
523  mtInv+G4      15374.124    544 31836.248    32041.354    35177.530
524  mtInv+I+G4    15367.084    545 31824.168    32030.099    35171.593
525  mtInv+R2      15349.539    545 31789.079    31995.009    35136.503
526  mtInv+R3      15332.068    547 31758.137    31965.724    35117.845
527  mtInv+R4      15324.469    549 31746.938    31956.189    35118.931
534  mtInv+F       15424.142    562 31972.285    32192.547    35424.124
535  mtInv+F+I     15315.536    563 31757.072    31978.194    35215.053
536  mtInv+F+G4    15304.048    563 31734.096    31955.218    35192.077
537  mtInv+F+I+G4  15296.945    564 31721.890    31943.876    35186.014
538  mtInv+F+R2    15278.574    564 31685.147    31907.133    35149.271
539  mtInv+F+R3    15261.456    566 31654.911    31878.628    35131.319
540  mtInv+F+R4    15253.972    568 31643.944    31869.401    35132.636
Akaike Information Criterion:           mtMet+F+R4
Corrected Akaike Information Criterion: mtMet+F+R4
Bayesian Information Criterion:         mtMet+F+R3
Best-fit model: mtMet+F+R3 chosen according to BIC

All model information printed to FINAL.fasta.model.gz
CPU time for ModelFinder: 2236.970 seconds (0h:37m:16s)
Wall-clock time for ModelFinder: 452.858 seconds (0h:7m:32s)
Generating 1000 samples for ultrafast bootstrap (seed: 940353)...

NOTE: 56 MB RAM (0 GB) is required!
Measuring multi-threading efficiency up to 48 CPU cores
Increase to 10 rounds for branch lengths
104 trees examined
Threads: 1 / Time: 48.189 sec / Speedup: 1.000 / Efficiency: 100% / LogL: -15327
Threads: 2 / Time: 26.375 sec / Speedup: 1.827 / Efficiency: 91% / LogL: -15327
Threads: 3 / Time: 19.241 sec / Speedup: 2.505 / Efficiency: 83% / LogL: -15327
Threads: 4 / Time: 15.826 sec / Speedup: 3.045 / Efficiency: 76% / LogL: -15327
Threads: 5 / Time: 14.712 sec / Speedup: 3.276 / Efficiency: 66% / LogL: -15327
Threads: 6 / Time: 13.501 sec / Speedup: 3.569 / Efficiency: 59% / LogL: -15327
Threads: 7 / Time: 12.408 sec / Speedup: 3.884 / Efficiency: 55% / LogL: -15327
Threads: 8 / Time: 11.746 sec / Speedup: 4.103 / Efficiency: 51% / LogL: -15327
Threads: 9 / Time: 11.425 sec / Speedup: 4.218 / Efficiency: 47% / LogL: -15327
BEST NUMBER OF THREADS: 8

Estimate model parameters (epsilon = 0.100)
1. Initial log-likelihood: -15311.614
2. Current log-likelihood: -15298.099
3. Current log-likelihood: -15286.477
4. Current log-likelihood: -15276.953
5. Current log-likelihood: -15269.839
6. Current log-likelihood: -15264.915
7. Current log-likelihood: -15261.662
8. Current log-likelihood: -15259.449
9. Current log-likelihood: -15257.634
10. Current log-likelihood: -15255.901
11. Current log-likelihood: -15254.208
12. Current log-likelihood: -15252.574
13. Current log-likelihood: -15251.022
14. Current log-likelihood: -15249.569
15. Current log-likelihood: -15248.221
16. Current log-likelihood: -15246.977
17. Current log-likelihood: -15245.827
18. Current log-likelihood: -15244.788
19. Current log-likelihood: -15243.809
20. Current log-likelihood: -15242.889
21. Current log-likelihood: -15242.059
22. Current log-likelihood: -15241.608
23. Current log-likelihood: -15241.318
24. Current log-likelihood: -15241.104
25. Current log-likelihood: -15240.924
26. Current log-likelihood: -15240.762
27. Current log-likelihood: -15240.612
28. Current log-likelihood: -15240.470
29. Current log-likelihood: -15240.333
30. Current log-likelihood: -15240.201
31. Current log-likelihood: -15240.072
32. Current log-likelihood: -15239.944
33. Current log-likelihood: -15239.818
34. Current log-likelihood: -15239.692
35. Current log-likelihood: -15239.565
36. Current log-likelihood: -15239.436
37. Current log-likelihood: -15239.306
38. Current log-likelihood: -15239.173
39. Current log-likelihood: -15239.037
40. Current log-likelihood: -15238.898
41. Current log-likelihood: -15238.756
42. Current log-likelihood: -15238.609
43. Current log-likelihood: -15238.457
44. Current log-likelihood: -15238.300
45. Current log-likelihood: -15238.139
46. Current log-likelihood: -15237.973
47. Current log-likelihood: -15237.801
48. Current log-likelihood: -15237.624
49. Current log-likelihood: -15237.441
50. Current log-likelihood: -15237.252
51. Current log-likelihood: -15237.057
52. Current log-likelihood: -15236.857
53. Current log-likelihood: -15236.650
54. Current log-likelihood: -15236.437
55. Current log-likelihood: -15236.219
56. Current log-likelihood: -15235.994
57. Current log-likelihood: -15235.764
58. Current log-likelihood: -15235.528
59. Current log-likelihood: -15235.288
60. Current log-likelihood: -15235.042
61. Current log-likelihood: -15234.793
62. Current log-likelihood: -15234.539
63. Current log-likelihood: -15234.283
64. Current log-likelihood: -15234.024
65. Current log-likelihood: -15233.763
66. Current log-likelihood: -15233.502
67. Current log-likelihood: -15233.240
68. Current log-likelihood: -15232.980
69. Current log-likelihood: -15232.721
70. Current log-likelihood: -15232.465
71. Current log-likelihood: -15232.213
72. Current log-likelihood: -15231.965
73. Current log-likelihood: -15231.723
74. Current log-likelihood: -15231.488
75. Current log-likelihood: -15231.259
76. Current log-likelihood: -15231.038
77. Current log-likelihood: -15230.826
78. Current log-likelihood: -15230.623
79. Current log-likelihood: -15230.429
80. Current log-likelihood: -15230.246
81. Current log-likelihood: -15230.072
82. Current log-likelihood: -15229.910
83. Current log-likelihood: -15229.758
84. Current log-likelihood: -15229.616
85. Current log-likelihood: -15229.484
86. Current log-likelihood: -15229.362
87. Current log-likelihood: -15229.250
88. Current log-likelihood: -15229.147
Optimal log-likelihood: -15229.037
Site proportion and rates:  (0.869,0.486) (0.120,3.547) (0.011,14.126)
Parameters optimization took 88 rounds (18.408 sec)
--------------------------------------------------------------------
|             INITIALIZING CANDIDATE TREE SET                      |
--------------------------------------------------------------------
Generating 99 parsimony trees... 1.998 second
Computing log-likelihood of 7 initial trees ... 0.627 seconds
Current best score: -15229.037

Do NNI search on 8 best initial trees
Finish initializing candidate tree set (10)
Current best tree score: -15229.037 / CPU time: 3.203
Number of iterations: 8
--------------------------------------------------------------------
|               OPTIMIZING CANDIDATE TREE SET                      |
--------------------------------------------------------------------
Iteration 10 / LogL: -15230.499 / Time: 0h:3m:15s
Iteration 20 / LogL: -15229.048 / Time: 0h:3m:15s
Iteration 30 / LogL: -15236.033 / Time: 0h:3m:16s (0h:7m:54s left)
Iteration 40 / LogL: -15229.038 / Time: 0h:3m:16s (0h:5m:3s left)
Iteration 50 / LogL: -15230.499 / Time: 0h:3m:17s (0h:3m:21s left)
Log-likelihood cutoff on original alignment: -15247.123
Iteration 60 / LogL: -15230.499 / Time: 0h:3m:18s (0h:2m:14s left)
Iteration 70 / LogL: -15230.499 / Time: 0h:3m:19s (0h:1m:26s left)
Iteration 80 / LogL: -15230.499 / Time: 0h:3m:20s (0h:0m:50s left)
Iteration 90 / LogL: -15229.038 / Time: 0h:3m:20s (0h:0m:22s left)
Iteration 100 / LogL: -15229.037 / Time: 0h:3m:21s (0h:0m:0s left)
Log-likelihood cutoff on original alignment: -15247.494
NOTE: Bootstrap correlation coefficient of split occurrence frequencies: 0.999
TREE SEARCH COMPLETED AFTER 101 ITERATIONS / Time: 0h:3m:22s

--------------------------------------------------------------------
|                    FINALIZING TREE SEARCH                        |
--------------------------------------------------------------------
Performs final model parameters optimization
Estimate model parameters (epsilon = 0.010)
1. Initial log-likelihood: -15229.037
2. Current log-likelihood: -15228.968
3. Current log-likelihood: -15228.923
4. Current log-likelihood: -15228.889
5. Current log-likelihood: -15228.865
6. Current log-likelihood: -15228.848
7. Current log-likelihood: -15228.835
Optimal log-likelihood: -15228.826
Site proportion and rates:  (0.879,0.491) (0.111,3.811) (0.010,14.283)
Parameters optimization took 7 rounds (1.517 sec)
BEST SCORE FOUND : -15228.826
Creating bootstrap support values...
Split supports printed to NEXUS file FINAL.fasta.splits.nex
Total tree length: 0.272

Total number of iterations: 101
CPU time used for tree search: 62.929 sec (0h:1m:2s)
Wall-clock time used for tree search: 10.430 sec (0h:0m:10s)
Total CPU time used: 880.940 sec (0h:14m:40s)
Total wall-clock time used: 204.399 sec (0h:3m:24s)

Computing bootstrap consensus tree...
Reading input file FINAL.fasta.splits.nex...
273 taxa and 547 splits.
Consensus tree written to FINAL.fasta.contree
Reading input trees file FINAL.fasta.contree
Log-likelihood of consensus tree: -15230.242

Analysis results written to: 
  IQ-TREE report:                FINAL.fasta.iqtree
  Maximum-likelihood tree:       FINAL.fasta.treefile

Ultrafast bootstrap approximation results written to:
  Split support values:          FINAL.fasta.splits.nex
  Consensus tree:                FINAL.fasta.contree
  Screen log file:               FINAL.fasta.log

Date and Time: Wed Mar 19 01:32:47 2025
